# Dynamic reinforcement learning reveals time-dependent shifts in strategy during reward learning

**DOI:** 10.1101/2024.02.28.582617

**Authors:** Sarah Jo C Venditto, Kevin J Miller, Carlos D Brody, Nathaniel D Daw

## Abstract

Different brain systems have been hypothesized to subserve multiple “experts” that compete to generate behavior. In reinforcement learning, two general processes, one model-free (MF) and one model-based (MB), are often modeled as a mixture of agents (MoA) and hypothesized to capture differences between automaticity vs. deliberation. However, shifts in strategy cannot be captured by a static MoA. To investigate such dynamics, we present the mixture-of-agents hidden Markov model (MoA-HMM), which simultaneously learns inferred action values from a set of agents and the temporal dynamics of underlying “hidden” states that capture shifts in agent contributions over time. Applying this model to a multi-step, reward-guided task in rats reveals a progression of within-session strategies: a shift from initial MB exploration to MB exploitation, and finally to reduced engagement. The inferred states predict changes in both response time and OFC neural encoding during the task, suggesting that these states are capturing real shifts in dynamics.

## Introduction

Behavior is rarely static. Time-varying factors, both internal and external, can influence the way in which humans and animals make decisions. Different ways of choosing an action can be attributed to using different strategies. One prominent perspective on such strategy heterogeneity is that the brain contains relatively independent, separable circuits that are conceptualized as supporting distinct strategies, each potentially competing for control. For instance, instrumental behavior appears to be supported by separable circuits for automatic versus deliberative control (***Killcross and Coutureau, 2003; Daw et al., 2005; Gremel and Costa, 2013***), and perhaps also different systems mediate the balance between such instrumental exploitation vs. different exploration strategies (***Daw et al., 2006; Blanchard and Gershman, 2018; Hogeveen et al., 2022; Cinotti et al., 2019; Cockburn et al., 2022***). The relative contribution of different strategies to behavior is often characterized via a “mixture-of-agents” (MoA) model that describes behavior as a weighted contribution of multiple “agents” that implement different strategies (***Camerer and Ho, 1999; Daw et al., 2011; Miller et al., 2017; Krueger et al., 2017; Gershman, 2018; Wilson et al., 2021; Cockburn et al., 2022***). It is hypothesized that the brain acts in the same way, weighing advice from separable circuits on how best to act (***O’Doherty et al., 2021***). The agent with the biggest weight can be thought of as the dominant strategy. Henceforth, we will broaden our use of the word “strategy” to now refer to a given weighting over the underlying individual agents.

Although one key goal of this type of modeling is to characterize how humans and animals switch between different strategies (e.g., the formation of habits with excessive training), in practice MoA models are usually fit to behavior assuming a single, fixed weighting over agents. Such models will therefore fail to capture changes in strategy over time, limiting our ability to associate behavioral and neural factors underlying variable strategies, and potentially missing contributions of a lesser used strategy all-together. For example, competing systems of automation and deliberation are often formalized and measured through a reinforcement learning (RL) framework as a mixture of model-free (MF) and model-based (MB) control (***Daw et al., 2005; Sutton and Barto, 2020***). A multi-step reward-guided task known as the “two-step task” was developed to differentiate these strategies in humans (***Daw et al., 2011***) and has since been used in rodents (***Miller et al., 2017; Hasz and Redish, 2018; Groman et al., 2019b; Akam et al., 2021***). Many versions of the task show choices are well fit by a mixture of both MB and MF behavior (***Daw et al., 2011; Hasz and Redish, 2018; Groman et al., 2019b; Akam et al., 2021***), whereas other studies show behavior dominated by MB planning (***Miller et al., 2017; Feher da Silva and Hare, 2020; Miller et al., 2022***). The lack of evidence for MF influences in these studies is somewhat puzzling, for instance because MF learning is predicted by classic models of the dopamine system (***Schultz et al., 1997***).

Apart from a few studies which build in some specific hypothesized change rule for strategy weighting (but do not, accordingly, measure such change in an unbiased way) (***Lee et al., 2014; Kool et al., 2016; Ebitz et al., 2018; Costa et al., 2019***), these studies neglect the dynamic representation of strategy. This leaves open the possibility that characterizing the dynamics of the agent mixture could reveal time-dependent trade-offs between strategies. For example, it might help clarify the behavioral discrepancies between different two-step tasks by revealing bouts of trials where MF control is the dominant agent in animals whose behavior is otherwise dominated by a MB agent. It might also identify dynamic shifts in the contributions from other less well understood agents, such as a novelty preference agent. This in turn might help to clarify its function, e.g. (as we show here) for exploration.

To answer these questions, we present an elaborated model that not only measures the contribution of separable agents, but also characterizes how this contribution changes over time. A recent class of models known as GLM-HMMs (***Calhoun et al., 2019; Ashwood et al., 2022; Bolkan et al., 2022; Oostland et al., 2022***) seeks to measure time-dependent shifts in strategy by nesting a generalized linear model (GLM), which maps perceptual variables to choices, under a hidden Markov model (HMM), which models an internal “hidden” state that switches between different GLM instantiations, producing a dynamic weighting over perceptual variables. This model has been successful at capturing how the dependence of actions on measurable variables — such as choice history, reward, or a task stimulus — can vary dynamically throughout a task. This can infer, for example, periods of engagement and disengagement (***Ashwood et al., 2020***), behavioral differences from neural inhibition (***Bolkan et al., 2022***), or speed of task learning with respect to an experimental stimulus (***Oostland et al., 2022***). However, GLM-HMMs, as originally specified, model dynamics over directly observable variables in the GLM and do not incorporate learning rules that generate internal variables themselves (such as action values inferred from models of reinforcement learning) that intervene to explain observed data.

Here we replace the GLM stage of the GLM-HMM with a MoA to produce the mixture-of-agents hidden Markov model (MoA-HMM). Extending both MoA models and GLM-HMMs, the MoA-HMM simultaneously characterizes the weighted influence of various inferred learning rules (and optionally observed variables) on subsequent actions and infers how this weighting over agents can dynamically shift over time. We apply this approach to reanalyze data from rats performing the two-step task (***Miller et al., 2017***) to examine whether there is evidence for strategy shifting, and if so — making no assumptions other than the HMM dynamics — how it proceeds. Using MB and MF learning rules to capture MB versus MF control alongside differentiating between exploratory and exploitative behavior, we show that the MoA-HMM reveals discernible shifts in the dominant agent throughout behavioral sessions. These shifts reveal a time-dependent trade-off between exploration and exploitation as well as drifting engagement in the task. We further show that these shifts in strategy provide a new lens to analyze behavioral features and neural data that were not used to train the model: the model-inferred strategy captures, out of sample, significant changes in both reaction times and neural encoding.

## Results

### Reinforcement learning in the rat two-step task

We study rat behavior and neural recordings in a multi-step, reward-guided decision-making task known as the rat two-step task. Originally described by ***Miller et al. (2017***), this task is an adaptation of a similar task in humans (***Daw et al., 2011***) that was designed to study the relative contribution of MB and MF strategies to choices. In the current version of the task, the rat is placed in an environment with two choice states (step one) that are separated in space and time from two outcome states (step two), in addition to initiation states for each step (**Figure 1A**). Each state is characterized by its own nose port, into which the rat has been trained to only poke when lit (yellow sun). The rat initiates a trial by entering the top center port (**i**), after which it is presented with a free decision between the two choice states (**ii**). Once a choice is made, the animal initiates the second step (**iii**) and is presented with a single outcome port (**v**) that it can enter for a chance at receiving liquid reward (**vi**). The available outcome port is dependent on a set, probabilistic link between choices in step one and outcomes in step two (**iv**), which can be represented as a transition matrix — the environment’s action-outcome model. Here one choice has an 80% probability of leading to one outcome (its *common* transition) and a 20% chance of leading to the opposite outcome (its *rare* transition), with flipped probabilities for the opposite choice. Either outcome state has a chance of giving reward; however, one outcome will always have a greater chance than the other (80% vs 20%), where the better outcome state reverses unpredictably throughout the session (**vii**). The goal of the rat, who’s been incentivized for liquid reward, is to track the better outcome port to maximize reward, which they do successfully (***Miller et al., 2017***).

**Figure 1.**
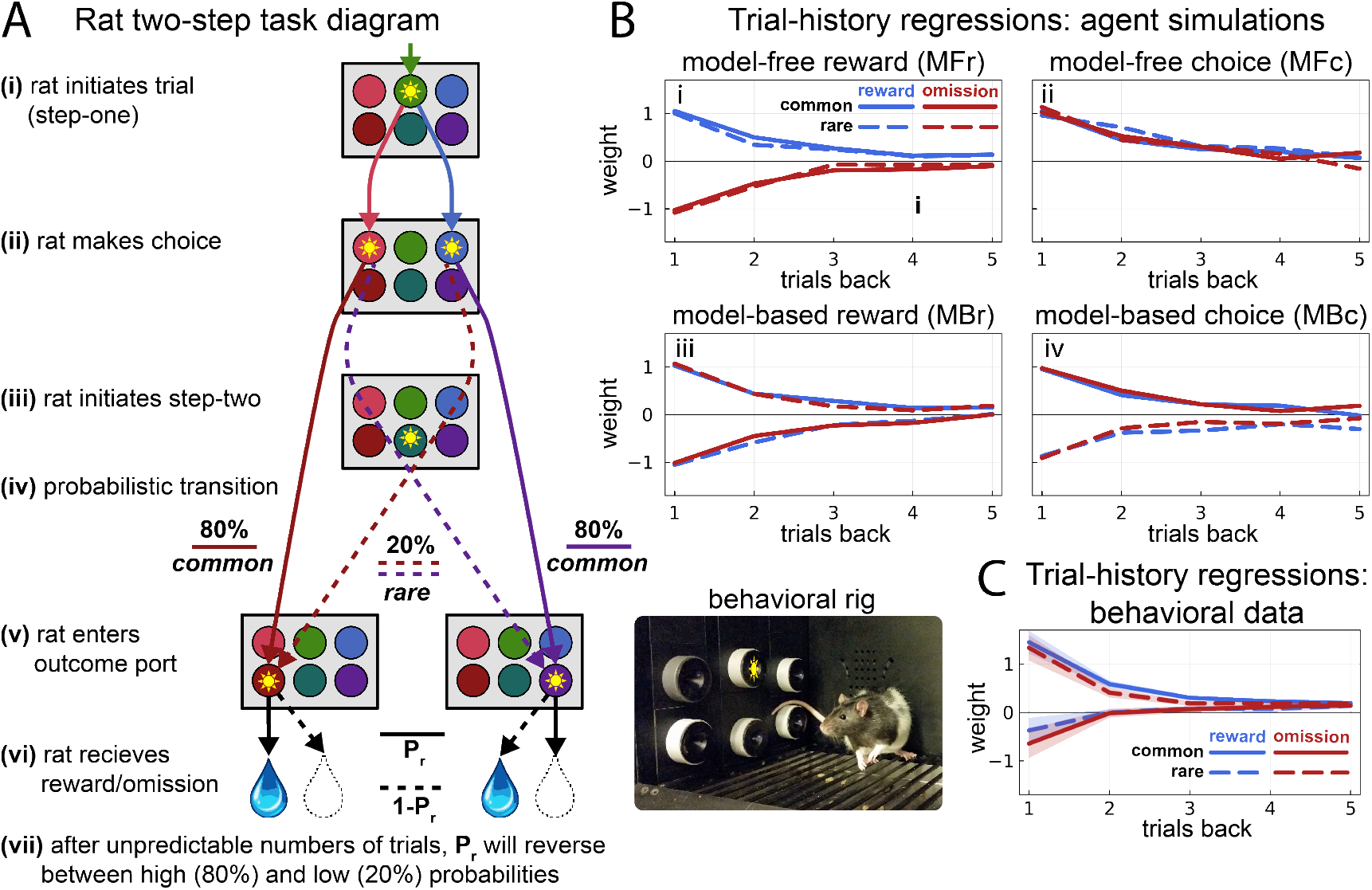
Rat Two-Step Task Description and Behavior. **A** Schematic of single trial during the rat two step task, where yellow suns indicate active port. The rat initiates a trial by poking into the top-center port (**i**), after which the rat is prompted to chose between the adjacent side ports (**ii**). After making a choice, the rat initiates the second step (**iii**). During the second step, the probabilistic transition determines the outcome probability (**iv**), and the rat is instructed to enter the active outcome port (**v**). One outcome port has some probability of delivering liquid reward, *P*_*r*_, and the other outcome port delivers reward with the inverse probability, 1 – *P*_*r*_ (**vi**). Throughout the session, the reward probability *P*_*r*_ will reverse unpredictably (**vii**). **B** Trial-history regressions fit to data simulated by each individual RL agent, each with learning rate 0.5 (5000 trials split between 20 sessions for each simulation). Each agent demonstrates differential effects of trial type on choices, which decays across past trials. (**i**) The model-free reward (MFr) agent tends to repeat choices (positive weight) after rewarded (blue) trials and switch away from choices (negative weight) following omission trials (red), ignoring the observed transition. (**ii**) Positive model-free choice (MFc) captures choice perseveration, tending to repeat choices regardless of observed reward or transition. (**iii**) The model-based reward (MBr) agent tends to repeat choices following common-rewarded (blue-solid) and rare-omission (red-dashed) trials, and switches choices following rare-rewarded (blue-dashed) and common-omission (red-solid) trials, showing an effect by both transition and reward. (**iv**) Model-based choice (MBc) agent with positive weight tends to repeat choices following common transitions (solid) and switch away from choices following rare transition trials (dashed), capturing transition-dependent outcome port perseveration. **C** Trial-history regressions fit to animal behavioral data (n=20 rats), where dark lines are mean regression values across all rats and shaded error bars are 95% confidence intervals around the mean. Rats look most similar to the MBr agent, but show additional modulation that can be accounted for by the influence of other agents.

The strength of the two-step task is in its separation of choice and outcome states by its transition structure, which allows MF and MB strategies to be distinguished by the way rewards and outcomes influence subsequent choices (**Figure 1B**). The key insight is that the stochastic state transitions (common vs. rare) decouple each choice’s reward history (which should drive MF learning) from the reward histories associated with its predominant outcome state (which could, alternatively, be used for MB evaluation of the choice’s value in terms of where it is likely to lead).

We can summarize MB and MF reward-learning strategies via two learning rules (or agents) that differ in whether they take account of transition in learning from reward. We also accompany them by two additional agents that capture reward-independent choice autocorrelation (perseveration or alternation), either MF (i.e., over choices) or MB (i.e., over outcomes or predominant destination states). These learning rules (based on the formulation from ***Ito and Doya, 2009***, extended to the two-step task, see ***Park et al., 2017; Groman et al., 2019a***) have some algebraic differences from variants used in some previous studies (***Daw et al., 2011; Miller et al., 2017***; see Methods), but capture the same logic. In addition to the four RL agents, an intercept or “bias” agent is included that captures the overall tendency towards left-ward or right-ward choices.

To demonstrate how reward and transition differently affect each agent, we adopt a trial-history regression that maps the influence of each trial type (common-reward, common-omission, rare-reward, and rare-omission) at varying trial lags (***Miller et al., 2016***). A positive weight indicates that choices tend to repeat following that trial type, and a negative weight indicate that choices tend to switch following that trial type. Specifically, the first agent, model-free reward learning (MFr) integrates choice and reward information by positively influencing choices that lead to reward and negatively influencing choices that lead to omissions (**Figure 1Bi**), ignoring the transition (outcome state) that was experienced. The second agent, model-free choice learning (MFc), uses only choice information and captures a history-dependent tendency to repeat or avoid choices on subsequent trials, i.e., *choice perseveration* or *alternation* (**Figure 1Bii**). In other words, depending on the sign of the agent’s mixture coefficient, each choice will favor or disfavor the same choice on future trials regardless of the reward received. Notably, a negative coefficient will drive a tendency to select the choice not previously sampled, akin to directed exploration in choice space. Put together, these agents can capture *MF exploitation* from a positive influence of MFr and *MF exploration* from a negative influence of MFc.

The third agent, model-based reward learning (MBr), captures the effect of both reward and transition on future choices (**Figure 1Biii**). The key feature is that a rare transition, relative to a common one, reverses the effect of a reward or omission on MB evaluation. This is because of the symmetry of the transition matrix: the reward received after a rare transition is at an outcome port which is most often reached by the other choice. Thus, rewards following common transitions and omissions following rare transitions will positively weight the selected choice, and omissions following a common transition and a reward following a rare transition will negatively influence the selected choice. The fourth agent, model-based choice learning (MBc), captures a history-dependent effect of the sampled outcome port on choices (**Figure 1Biv**). Put simply, it captures a “common-stay, rare-switch” pattern of choices during the two-step task, leading to *outcome perseveration* or (if negative) *alternation*. That is, like MFc, this agent tends to repeat or switch choices regardless of reward; however, it does so by choosing or avoiding the choice commonly associated with the obtained outcome port rather than the choice itself. This means that (if negative) it promotes switching choices so as to sample previously unsampled outcomes, like a MB (transition-sensitive) form of directed exploration. This comes from repeating choices after “rare” transitions in attempt to sample the “common” outcome, and switching choices after “common” transitions in attempt to sample the opposite “common” outcome. These agents, then, can capture *MB exploitation* from a positive influence of MBr and *MB exploration* from a negative influence of MBc.

As shown previously (***Miller et al., 2017***), the rats’ pattern of choices most closely follow the MBr agent (**Figure 1C**). It should be noted, however, that the RL agents used here differ from those previously used to model the rat two-step task (***Miller et al., 2017, 2022***). The main elaboration is the inclusion of MBc to capture history-dependent outcome perseveration/MB exploration. This generalizes and clarifies the role of the “novelty preference” (NP) agent used in the earlier studies, which similarly captured “common-stay, rare-switch” behavior (albeit in the opposite direction, i.e. “rare-stay/common-switch”), but lacked dependence on more than the preceding trial. Although it significantly contributed to behavior previously, its role was not thoroughly explored. The updated learning rules used here not only provide a better fit to behavior than previous versions of the MoA, (Figure 5 – figure supplement 1), but also the role of the MBc agent is further clarified especially, as we’ll see later on, when its dynamics become apparent. For further detail on specific value update rules for the new and original agents used for comparison, refer to the methods section.

### MoA-HMM framework

Before assessing how an MoA-HMM can describe behavior, it’s important to first understand how a static MoA model can generate decisions. Imagine a set of agents, *A* ∈ {*agents*}, and a set of choices, *y* ∈ {*choices*}, where each agent has its own value for each choice *Q*_*A*_(*y*) that is updated at the end of every trial. Using the term “actor” to refer to overall choice selection, a MoA model dictates that the probability of selecting a choice *y*_*t*_ on some trial *t* arises from a softmax over net valuations for each choice, computed as the weighted sum of each agent’s values

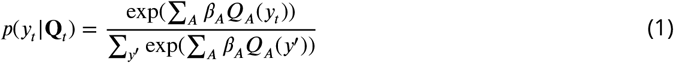

In the context of the two-step task (and similar bandit-style reward learning tasks more generally), an actor takes some choice *y* that leads to some outcome state *o* and reward *r*. For demonstration, we assume an actor’s behavior is described by an MoA with two agents, a model-free choice learning rule (MFc) and a model-based reward learning rule (MBr), *A* ∈ {MBr, MFc} (**Figure 2A**), which might reflect competition between goal-directed and habitual strategies (***Miller et al., 2019***). On the first trial, each agent’s values are initialized (e.g., uniform across choices), **Q**_1_ = [*Q*_MBr_, *Q*_MFc_] (**i**). A net value is computed as a weighted sum of **Q**_1_, given static agent weights ***β*** = [*β*_MBr_, *β*_MFc_], to produce a probability distribution over choices *p*(*y*_1_) (**eq. 1**), from which the actor’s choice is drawn **(ii)**. After a choice is made **(iii)**, some outcome *o*_1_ and reward *r*_1_ is observed by the actor **(iv)**, and all observed variables may be used to update each agent’s choice values **(v)**. These values are fed into the next trial, and this process repeats until the final trial.

**Figure 2.**
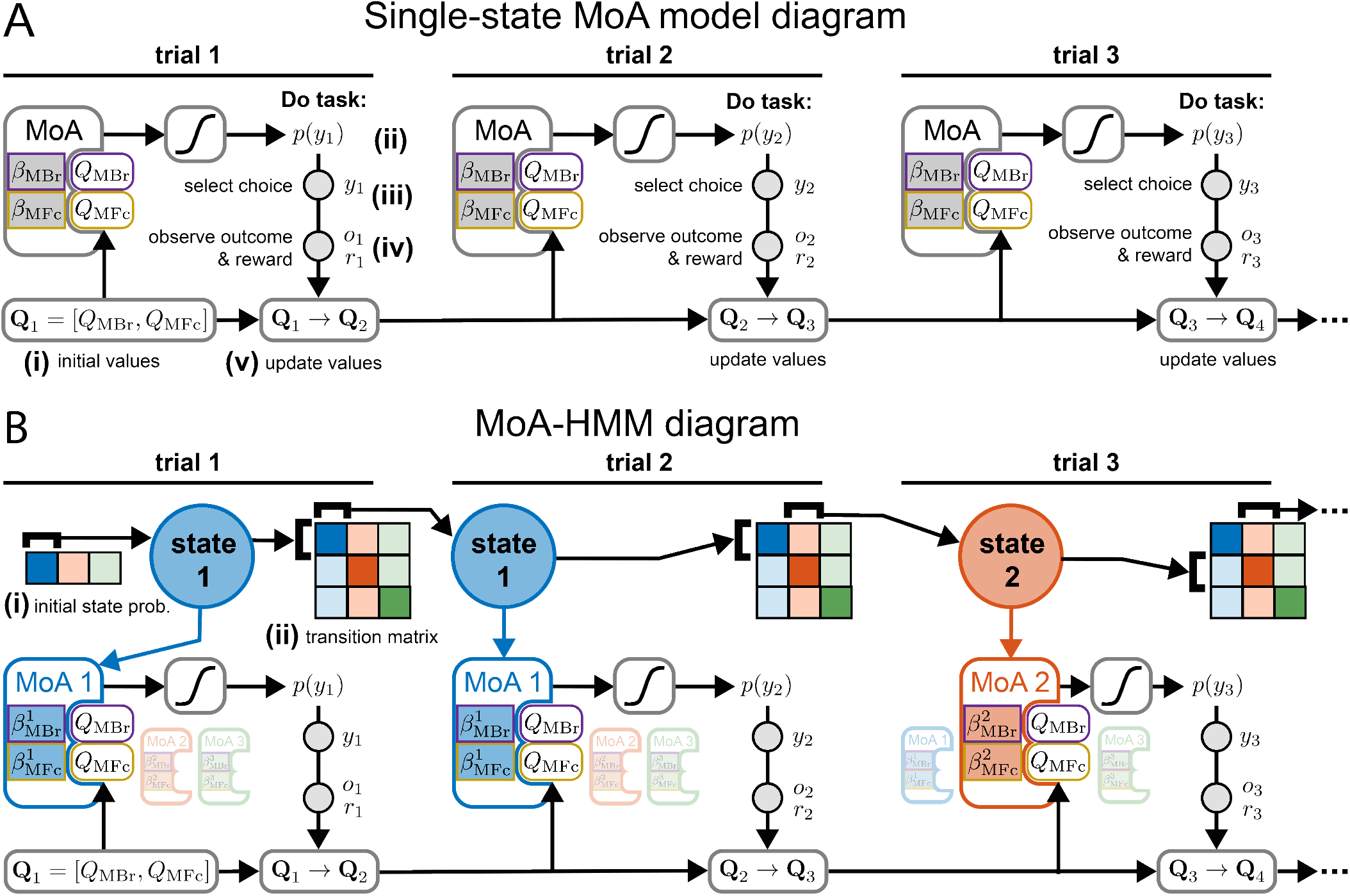
Model diagrams during a generic reward learning task. **A** Diagram of a static MoA model. At the start of a session, values for each agent, **Q**_1_ = [*Q*_*MBr*_, *Q*_*MF c*_], are initialized (**i**) and fed into a MoA described by agent weights ***β*** = [*β*_*MBr*_, *β*_*MF c*_]. This weighted sum is pushed through a softmax to produce a probability distribution *P* (*y*_1_) (**ii**) from which to draw choice *y*_1_ (**iii**). An outcome port *o*_1_ and reward *r*_1_ are observed (**iv**), and all task variables may be used to update agent values according to their respective update rules (**v**). These updated values are used by the MoA on the next trial, and this process repeats until the session terminates. **B** Diagram of a MoA-HMM model. Values are initialized and updated in the same manner as (A), but now the MoA used to generate choice probability is determined by an underlying hidden state, where each hidden state has its own set of agent weights ***β***^*z*^ = [***β***^1^, ***β***^2^, ***β***^3^]. The state at the beginning of the session is determined by an initial state probability distribution (**i**), and the conditional probability of each subsequent hidden state is governed by a transition matrix (**ii**), each row corresponding to the previous state and each column corresponding to the upcoming state.

Next consider an HMM in isolation. An HMM produces a stochastic trajectory of hidden states that are each associated with different probabilistic patterns over observable data. Thus, it produces discrete changes in observation statistics, reflecting the underlying, temporal transition dynamics of hidden states. When paired with an MoA, these hidden states determine the set of weights being used to combine agents at each time step (the *β*s in Equation 1), and thereby imply dynamics in how these sets of weights trade off over time. We assume that the MoA-HMM uses the same choice values and learning rules as before, but the weighting over agent values is now dependent on hidden state 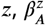. Thus, at time point *t* given hidden state *z*_*t*_, equation 1 updates to

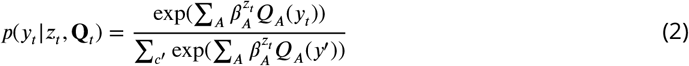

**Figure 2B** demonstrates how an MoA-HMM with three hidden states affects how choices are generated from two RL agents in an MoA model. Notably, in the case of three hidden states, we now have three sets of weights ***β***^*z*^ = [***β***^1^, ***β***^2^, ***β***^3^], where the set is selected according to the active hidden state. The active hidden state at any given time is, in turn, influenced by two distributions governing the HMM dynamics: 1) the initial state distribution, *p*(*z*_1_), determines the state at the beginning of a sequence **(i)**; and 2) the conditional probability of a subsequent state given the current state *p*(*z*_*t*_ | *z*_*t*−1_), i.e. a state transition matrix **(ii)**, iteratively determines each subsequent state in the sequence. (See methods for more details on the fitting procedure.)

How might an MoA-HMM improve our understanding of RL behavior during a reward-learning task? As stated previously, RL behavior is often modeled using a weighted mixture of model-free and model-based RL rules. This simple mixture model would fail fully to capture a situation in which one learning rule dominates during some blocks of trials and the other dominates during other blocks of trials. For example, consider an actor during the two-step task whose behavior can be described by three states, one that is more goal-directed in which MBr dominates with a slower learning rate, a second that is more habitual in which MFc dominates with a faster learning rate, and a third producing a heavy side bias (**Figure 3Ai,ii**). Assume this actor also has a non-uniform initial-state probability (**iii**) and asymmetric transition dynamics (**iv**). Accordingly, over the course of a session, the net values used to select choices would be primarily dominated by one of the agents dependent on which hidden state is active (**Figure 3B**), allowing for variability, clustered in time, in how these values contribute to choice selection. Fitting a single-state MoA model to data generated from this actor would describe behavior as an even mix of MBr and MFc agents (**Figure 3Ci**) and might even fail to recover the MBr and MFc learning rates (**ii**). This single-state MoA would be unable to capture the dynamic influence of each agent with no estimation of the underlying hidden state (**Figure 3D**). A 3-state MoA-HMM, however, is able to recover the original model parameters (**Figure 3E**) as well as approximate the underlying hidden state and dynamic influence of agents (**Figure 3F**). Beyond this didactic example, 3-state MoA-HMMs can be well-recovered over a wide range of parameters even when including all four RL agents (Figure 3 – figure supplement 1 for recovery of randomized parameters with noted limitations, and figure supplement 2 for recovery of models fit to real data). To test if it can reveal switching RL strategies in real data, we apply the MoA-HMM to rats behaving in the two-step task.

**Figure 3.**
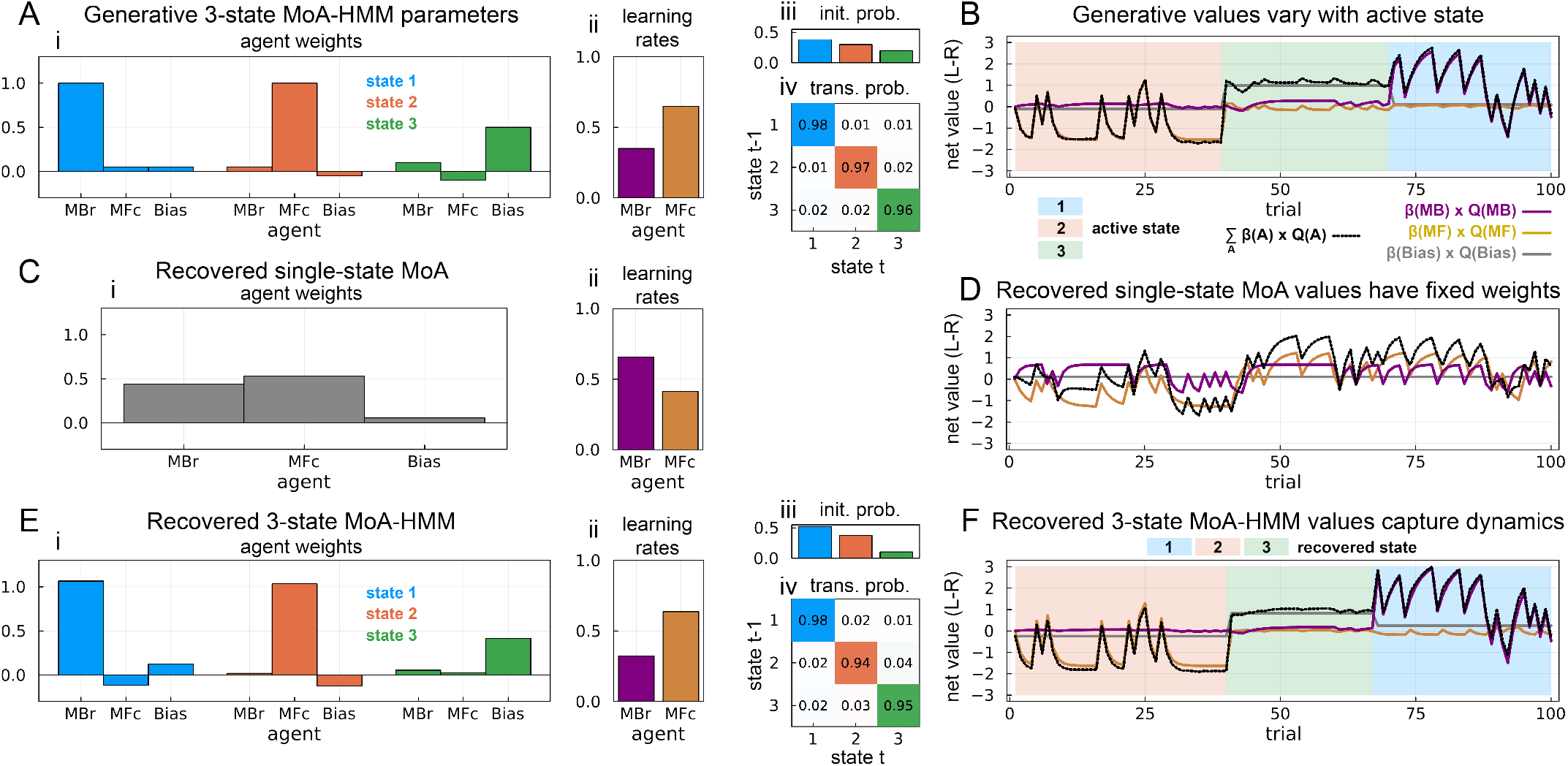
Example 3-state MoA-HMM with single-state and 3-state model recovery. **A** 3-state MoA-HMM parameters used to generate example data during the two-step task. Three agents are included: MBr, MFc, and Bias. (**i**) Agent weights for each hidden state. The first state (blue) is dominated by the MBr agent, the second state (orange) is dominated by the MFc agent, and the third state (green) is dominated by the Bias agent. (**ii**) Learning rates for MBr and MFc agents. (**iii**) Non-uniform initial state probability. (**iv**) Asymmetric transition probability matrix. **B** Effective (weighted) agent values during an example session. The dominant value in the weighted sum (black line) changes depending on the active hidden state (background color). **C** Recovered parameters by a single-state MoA. (**i**) Recovered agent weights erroneously identify behavior as a mixture of the three agents. (**ii**) Agent learning rates recovered by the single-state MoA. **D** Inferred values from recovered single-state MoA. The weighting on each agent is fixed across the session. **E** Recovered 3-state MoA-HMM parameters. (**i**) Each generative state, with a single dominating strategy per state, is recaptured. (**ii**) Recovered learning rates. (**iii**) Recovered initial state probability. (**iv**) Recovered transition probability matrix. **F** Inferred values from recovered 3-state MoA-HMM vary similarly to the generative values. The recovered hidden state is also closely approximated. **Figure 3—figure supplement 1**. Simulated 3-state MoA-HMM parameter recovery for the five agents used in behavioral fits: model-based reward, model-based choice, model-free reward, model-free choice, and bias. Simulated model parameters were randomly drawn (see methods). Each simulation contained 5000 trials evenly split between 20 sessions. Parameters in each panel are pooled across states. “r” is the Pearson’s correlation coefficient between the simulated and recovered parameters, and “R2” is the coefficient of determination, *R*^2^, calculating how well the simulated parameters predict the recovered parameters. Due to the interaction between different model parameters (e.g. a small *β* weight will affect the recoverability of the agent’s learning rate *α*), a number of “failures” can be seen. **Figure 3—figure supplement 2**. 3-state MoA-HMM parameter recovery from data simulated from each rats’ behavioral model fit. Each rat’s model (n=20) was used to generate 5 independent data sets, where each data set contained the same number of trials and sessions as the corresponding rat’s behavioral data set used to fit the behavioral model, giving a total of 100 simulations. “r” is the Pearson’s correlation coefficient between the simulated and recovered parameters, and “R2” is the coefficient of determination, *R*^2^, calculating how well the simulated parameters predict the recovered parameters. Parameters inferred from real data are more reliably recovered.

### Dynamic reinforcement learning behavior is better captured by the MoA-HMM

We return to using a MoA model with all four RL agents alongside a “bias” agent. As shown previously (***Miller et al., 2017***), the rat two-step task specifically isolates MB planning in rats. Alongside the choice plot in Figure 1C, this can be further demonstrated by fitting a single-state MoA (to each rat’s data separately) and looking at the difference in normalized likelihood (cross-validated across sessions) when removing one of the agents (**Figure 4A**, gray). The normalized likelihood (computed as *exp*(*L*/*N*), where *L* is the cross-validated log likelihood of the model and *N* is the total number of trials) represents the proportion of the model likelihood attributed to the selected choice, where 50% is the model performing at chance level. We see the largest reduction in normalized likelihood when removing MBr (median of −3.10% across rats, p<1E-5 from a non-parametric signed rank test) and a non-significant change when removing MFr (median of −0.03%, p=0.08), consistent with results from ***Miller et al. (2017***). We also see significant decrease in likelihood when removing MBc (median of −0.34%, p<1E-4), MFc (median of −1.23%, p<1E-5), and Bias (median of −0.30%, p<1E-5 SR) agents.

**Figure 4.**
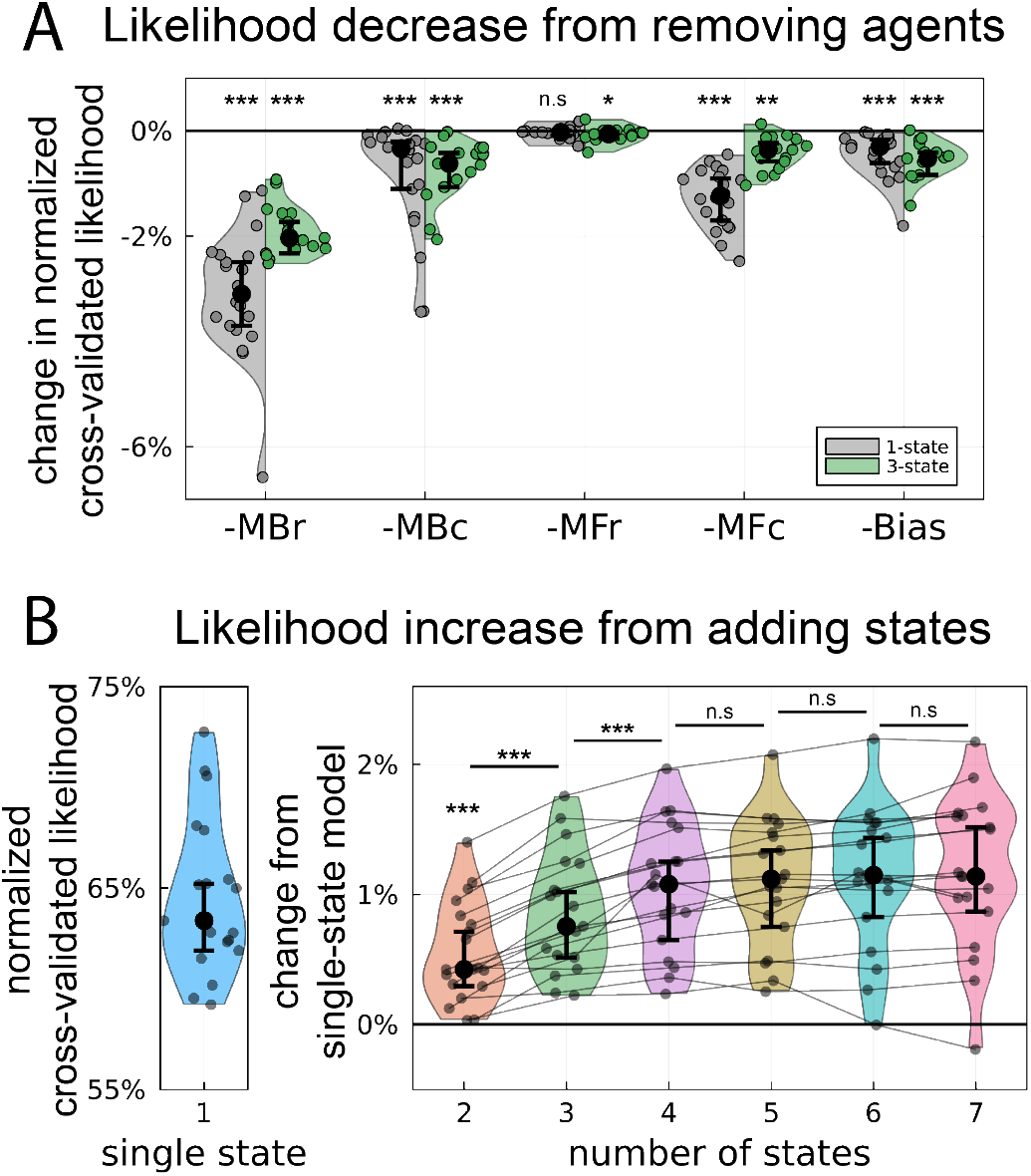
Changes in model fit quality across rats. **A** Change in normalized, cross-validated likelihood from the full model when removing one of the agents, computed for both a single-state MoA (gray, left) and a 3-state MoA-HMM (green, right). All agents, excluding MFr for a single-state MoA, show a significant (across rats) decrease in likelihood. **B** (left) Normalized, cross-validated likelihood of a single-state MoA. (right) Change in normalized, cross-validated likelihood when adding additional hidden states into the MoA-HMM, relative to the single-state model. Significant changes are computed with respect to models with one fewer states (e.g. 2-state vs 1-state, 3-state vs 2-state), where significant increases are observed through 4 states. **A-B** Each colored dot is an individual rat, black dots correspond to the median across rats, and error bars are bootstrapped 95% confidence intervals around the median. *:p<0.02, **:p<0.001, ***:p<1E-4 **Figure 4—figure supplement 1**. New RL learning rules significantly improve fit to behavior and capture much of the variance explained by the Novelty Preverence and Perseveration agents of the original model from ***Miller et al. (2017***). *:p<0.02, **:p<0.001, ***:p<1E-4

**Figure 5.**
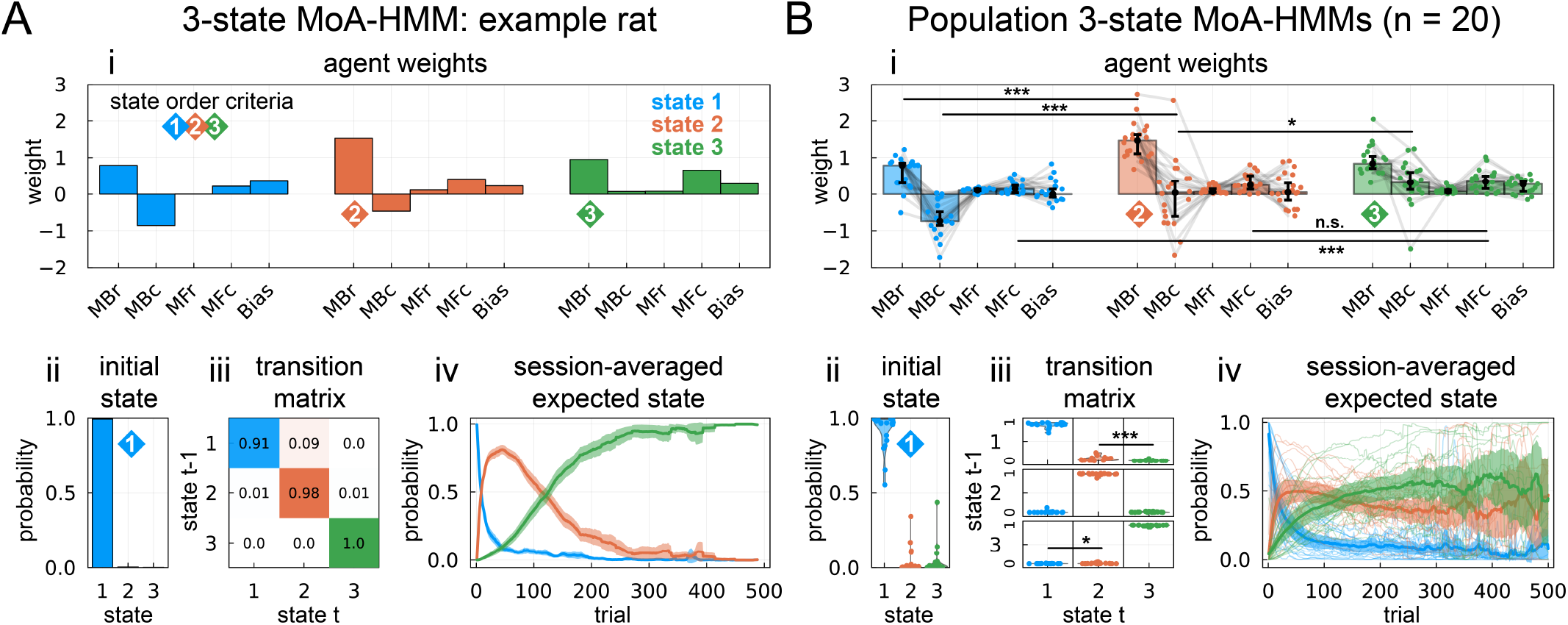
Example and summary 3-state MoA-HMMs fit to rats in the two-step task using a population prior (see methods). The states are manually ordered based on three properties: the first state is chosen as the one with the highest initial state probability (blue diamond), the remaining two states are then ordered second and third (orange and green diamonds, respectively) in order of the weight they give to MBr (orange = higher MBr). **A** Example 3-state MoA-HMM on a single rat. (**i**) Agent weights split by hidden state. (**ii**) Initial state probability. (**iii**) State transition probability matrix. (**iv**) The expected hidden state calculated from the forward-backward algorithm averaged (mean) across sessions, with error bars as 95% confidence intervals around the mean. **B** Summary of all 3-state MoA-HMM fits across the population of 20 rats. States were sorted in the same way as (A). (**i**) Agent weights split by hidden state, where individual dots represent a single rat, light grey lines connect weights within a rat, bars represent the median weight over rats, and error bars are bootstrapped 95% confidence intervals around the median. **(ii)**Distribution of initial state probabilities, with each dot as an individual rat. (**iii**) Distribution of state transition probabilities, with each panel representing a single (*t* − 1) → (*t*) transition. (**iv**) Session-averaged expected state computed as in (Aiv), where light lines are average probabilities for individual rats and the dark solid lines are the population mean with 95% confidence intervals around the mean. *:p<0.05, ***:p<E-4 **Figure 5—figure supplement 1**. (**A**) Learning rates fit for each agent (**i**) corresponding to the example rat shown in Figure 5A and (**ii**) summarizing each learning rate over the population of rats. Each dot is an individual rat, bars represent the median, and errorbars are bootstrapped 95% confidence intervals around the median. (**B**) Three example sessions showing the inferred state likelihood on each trial from the example rat shown in Figure 5A. (**C**) Cross-correlation between left choices and reward probabilities for the common outcome port given that choice (gray). Left choices are highly correlated to left-outcome reward blocks, with the peak correlation at a slight lag (vertical dashed line) indicating the trial at which the rat detects the reward probability flip. To test whether the latent states track reward flips, the cross correlation is also shown between left-outcome reward probability and the likelihood of each state: initial state (blue), the remaining state with a more rightward choice bias (orange), and the remaining state with a more leftward bias (green). These correspond directly to states 1-3 in the example rat (**i**) whose model is shown in Figure 5A. while other rats had states 2 and 3 assigned according to their individual choice biases. **Figure 5—figure supplement 2**. A second 3-state MoA-HMM example and comparison to a GLM-HMM. States identified by a MoA-HMM and GLM-HMM are highly similar. **A** Example 3-state MoA-HMM model parameters with (**i**) agent weights split by state, (**ii**) each agent’s learning rate, **(iii)** the initial state probability, and (**iv**) the state transition matrix. **B** 3-state GLM-HMM fit to the same rat as (A). (**i**) GLM-HMM regression weights for each state. Each state is described by four types of regressors indicating four possible trial types – common-reward, common-omission, rare-reward and rare-omission – and choice direction for up to 5 previous trials, giving 20 parameters per state. Each state additionally had a bias term (**ii**), leading to a total of 63 model weights for a 3-state model. (**iii**) GLM-HMM initial state probabilities also identify a prominent initial state. (**iv**) GLM-HMM transition matrix closely matches MoA-HMM. **C** Expected state probabilities for MoA-HMM (**i**) averaged across all sessions and (**ii**) highlighted example sessions. **D** Expected state probabilities for GLM-HMM (**i**) averaged across all sessions and (**ii**) highlighted example sessions. Temporal structure is highly similar to MoA-HMM. **E** Cross-correlation between the expected state probabilities inferred from the MoA-HMM and GLM-HMM (i.e. panels Cii and Dii) across all sessions. Each dot is an individual rat, black circles are medians and error bars are 95% confidence intervals around the median. **Figure 5—figure supplement 3**. Two example 4-state MoA-HMM fits corresponding to 3 state fits from (**A**) Figure 5A and (**B**) Figure 5-figure supplement 2. States are ordered according to the initial state probability (Aii and Bii) and the transition probabilities to most-likely states that follow (Aiii and Biii). Initial states are generally consistent with the 3-state fits, and the way the remaining two states split into three states is more idiosyncratic. For example, (**A**) suggests state 3 from the smaller model is split into two states (iv) that differ by bias (i), while (**B**) suggest the additional state 4 draws from both the smaller model’s states 2 and 3 (iv), and the state with largest MBr state no longer directly follows the initial state (i). computationally intensive.

What this model does not reveal, however, is any underlying dynamics to the dominant strategy. By adding additional hidden states, we observe a significant increase in normalized, cross-validated likelihood, suggesting that additional states capture dynamics in the behavior unaccounted for by a single-state MoA. (**Figure 4B**). At two hidden states (orange), we see an average jump of 0.77% (median, p<1E-5). The fit is improved further with three hidden states (green), seeing an median of 1.15% jump in quality vs the one-state model, which is also significantly greater than with two states (median increase of 0.33%, p<1E-4). Significant improvements can be seen until four states (purple, median of 1.37% vs one state; median increase of 0.18% vs three states, p<1E-4), after which we see non-significant gains from five or more states relative to the predecessor model (i.e. one fewer states). This suggests that four hidden states optimally describe behavior. However, our chief goal in what follows is to explore a descriptive model. Examination revealed that four states often highlight idiosyncrasies between animals (Figure 5 – figure supplement 3), making it more difficult to draw general conclusions. Therefore, in the remaining analyses we restrict our model to three states to facilitate interpretability and comparison across animals.

Using a three-state model, we find that the state dynamics absorb some of the variance when either the MBr or MFc are removed (**Figure 4A**, green, MBr: −2.03%, p<1E-4; MFc: −0.35%, p<0.001). Intriguingly, the removal of MBc and Bias both show, on average, a greater decrease (compared to a single-state model) in model fit likelihood (MBc: −0.62%, p<1E-4; Bias: −0.52%, p<1E-4) as well as MFr showing a slightly greater, now significant decrease in likelihood (−0.06%, p=0.015), possibly suggesting a greater role of these agents when dynamics are included.

### Time-varying hidden states reveal new dynamics of behavior

Fitting a three-state MoA-HMM to each rat reveals interesting properties not seen in a single-state MoA. **Figure 5A** shows an example fit on one of the animals, where we summarize the fit by displaying the agent weights split by hidden state (**i**), the initial state probability (**ii**), and the state transition probability matrix (**iii**). Additionally, we show the average likelihood of each state across sessions (**iv**), which displays the dynamics of when each state is inferred to be active within a session, averaged over sessions. From this example, we can identify some prominent features that are common to many of the rats. Since the ordering of states in the model is arbitrary, we used some of these features to identify a canonical ordering for the states across rats to allow group-level comparison, indicated by colored diamonds.

First, the initial state probability (**ii**) reveals that a single state dominates at the beginning of the session (blue diamond), which we’ve used to label the first state (blue). Comparing the agent weights (**i**) during this initial state to the remaining states, we notice that one weight in particular, MBc, is more negative than in the remaining two states. Recall that a negative weight for MBc captures a preference to visit outcome states not recently visited, which we identified as MB exploration. That this arises most strongly at the beginning of the session is consistent with this interpretation since the reward location is initially unknown. After the initial reward location is identified, exploratory behavior is less beneficial; with only two outcome states, there is only one alternate outcome once the other becomes less rewarding. This is corroborated by the decreasing magnitude/more positive weight on MBc in the later two states.

Since we want to identify changes in planning behavior, we selected the second state (orange) based on having the highest MBr weight (orange diamond) among the remaining states. In our example rat, this state also has a higher MBr weight than the initial state and is most likely to follow the initial state, seen in the transition matrix (**iii**) and average expected state probability (**iv**). Together with the reduced weight on MBc, this suggests a shift toward increased MB exploitation after an initial exploratory phase.

The remaining state is labeled green and, by definition, has a lower MBr weight than the second state (green diamond). Additionally, this state shows the most positive weight on both MBc and MFc, suggesting a stronger influence of perseveration (in both choices and outcomes) during this state. This state tends to occur towards the end of sessions; it is possible that later in a session, the rat is sated on water and is less motivated by reward-seeking. This is accompanied by an increase in less cognitively-demanding strategies such as perseveration, a pattern captured by the reduced magnitude on MBr and more positive weights on MBc and MFc.

Many of these properties hold across our population of rats (**Figure 5B**), coloring the states according to the criteria described above. One state (**ii**) tends to occur with high initial probability and, on average, the most negative weight on MBc (**i**, median of −0.74 in state 1; vs 0.04 in state 2, p<0.001; vs 0.32 in state 3, p<0.001 across rats). The second state, again selected by having the larger MBr weight of the remaining two, follows the initial state on average (**iii**, state 1 to 2 transition median: 0.037; vs state 1 to 3: 0.001, p=1E-4). Alongside the increase in MBc, the second state shows a significant increase in MBr from the first state (state 1 median: 0.78; vs state 2: 1.47, p<1E-5). The remaining state, defined by a decrease in MBr, is on average most active at the end of the session (**iv**). It sees the most positive MBc (median of 0.32; vs 0.04 in state 2, p=0.03) and MFc (median of 0.35; vs 0.15 in state 1, p<1E-4), although MFc is not significantly greater than state 2 (median of 0.25, p=0.35). Together with the lower MBr and higher MBc, however, the relative increase in MFc from the beginning of the session is consistent with the idea of reduced engagement and reduced motivation from reward.

Contrary to our initial expectation, we did not see any significant changes in MFr between states across our population of rats (state 1 median: 0.10; state 2 median: 0.06; state 3 median: 0.07). This further supports that rats are not substantially employing a MFr-like strategy during this version of the task, even during smaller subsets of trials. Thus, these data reveal no evidence of a trade-off between MBr and MFr strategies in this setting, though they do suggest one between alternation/perseveration strategies in choice vs outcome space (i.e., MBc vs. MFc).

To verify that these state dynamics are not an artifact of the particular agents as we have selected, we compared these results to states fit by a GLM-HMM using the same more generic, modelagnostic trial-history regressors as in Figure 1 (Figure 5 – figure supplement 2). Strikingly, the hidden state dynamics (i.e. initial state and transition matrix) are extraordinarily similar, and the expected state likelihoods from both models are highly correlated. The hidden states themselves reveal the same effects by reward and transition across both models; however, the MoA-HMM provides a more streamlined description of this behavior with far fewer parameters and more interpretable learning rules.

### Strategy shifts significantly predict changes in behavioral and neural metrics

#### Changes in strategy correlate to changes in response time

So far we have suggested an interpretation that might explain the progression of states we see across rats: an initial exploration state, a middle exploitation state, and an ending state with reduced engagement. To investigate this interpretation, we next looked to how the states relate, out of sample, to other measurements. If the relative influence of learning rules is changing between states, signifying a shift in strategy, we reasoned that response times may also change as different strategies may require different amounts of computation. For example, it is often thought that more MB strategies elicit slower response times than MF strategies, as MB strategies are more computationally intensive.

Our response time of interest is when the animal is most likely to engage in the learning rule updates we’re modeling; during the rat two-step task, this likely occurs during the inter-trial interval (ITI) following outcome receipt and prior to initiating the next trial (***Miller et al., 2022***) (**Figure 6i**). To understand whether latent state is related to ITI duration, we must also account for other variables that may also affect this response time. Thus, we account for the presence of reward due to time spent drinking. Additionally, since the progression of states is correlated with time within the session, we wish to ensure that the latent state per se is informative beyond any effects on response time that can simply be explained by the passage of time. To investigate this, we first transformed ITI duration to ITI rate (**Figure 6ii**) to constrain long ITIs and reveal a more linear relationship between ITI duration and time. We then fit a linear multilevel model to each rat with state identity, reward, and linear and nonlinear effects of time via a third degree polynomial (*time* + *time*^2^ + *time*^3^, together referred to as “time”) as predictors, as well as interactions between reward and both state and time, while capturing variance in all predictors due to session-level random effects.

**Figure 6.**
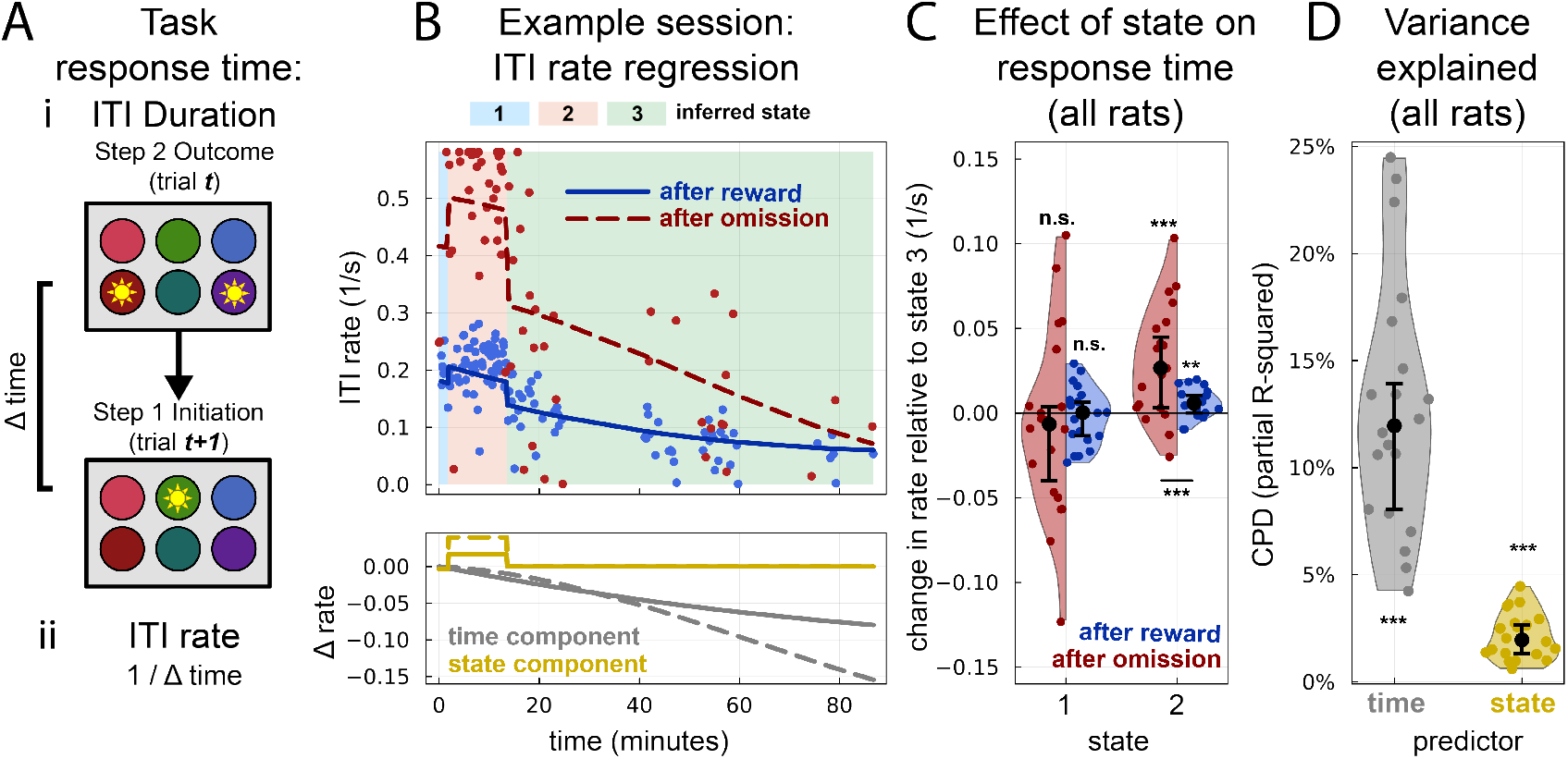
Changes in inter-trial interval (ITI) duration predicted by latent state. **A** Task diagram of response time of interest: ITI duration. (**i**) The ITI duration is measured as the difference in time (Δ*time*) between outcome port entry during step 2 of trial *t* and initiation port entry during step one of the next trial *t* + 1. (**ii**) Due to a large right tail from the longest ITIs, we instead model the inverse of ITI duration, or ITI rate, in our multilevel linear regression model. **B** Example prediction from a multilevel linear regression with fixed effects of reward, time, state, time × reward, and state × reward, and random effects from session on every predictor. (**top**) Scatter plot of ITI rate by time in session (in minutes) within an example session, colored by whether the ITI followed a reward (blue) or omission (red). Overlaid lines are the model-predicted rate split for reward (solid) and omission (dashed) trials. Discrete shifts in ITI rate align with shifts in inferred state (background color), with the highest rate during state 2. (**bottom**) Components of the prediction in the top panel due to time predictors (gray) and state predictors (gold) split for reward (solid) and omission (dashed) trials. The state component, and its interaction with reward, capture the discrete shift observed in the top panel. **C** Grouped dot plot of ITI rate split by inferred state, after subtracting changes in rate due to time (time component from i, bottom panel). State 2 shows a significantly higher rate than state 3 for ITIs following both rewarded and omission trials. **D** Coefficient of partial determination (CPD, or partial R-squared) for both time and state decouples the variance explained by each group of predictors. Time predicts significantly more than state, but state still significantly increases the variance explained by the model. **:p<0.01, ***:p<1E-4

**Figure 6B** shows an example regression predicted on one of the sessions. Strikingly discrete shifts in response times are visible, well aligned to the inferred state (**Figure 6Bi** top panel) and captured by the predictive component due to state identity (**Figure 6Bi** bottom panel). After subtracting out drifts in ITI rate due to time, we find significant differences in ITI rate split by inferred state (**Figure 6Bii**) in this example session, where state 2 sees a significantly higher rate (or shorter ITI duration) than state 3 following both rewarded and omission trials. This effect holds across rats, where fitting a multilevel model to each rat reveals a significant increase in ITI rate following omission (0.026 ITIs/s, p<0.001) and reward (0.006 ITIs/s, p=0.008), where the effect after omission is significantly greater (p<0.001). Since state 2 has the highest MBr weight, this may seem to contradict the expectation that MB strategies have slower response times than MF strategies. However, these states do not reflect a trade-off between MB and MF strategies; the fastest response time here is associated with the exploitative state, while slower response times are associated with the exploration and reduced engagement states. To formally disentangle the separable contributions of time and state, we use the coefficient of partial determination (CPD, or partial R-squared) to measure the explanatory power of each predictor. We find that while time contributes more to the model (median CPD=12%, p<1E-5, perhaps not surprisingly since it is, conservatively, modeled with a very flexible third-order form), the state identity still significantly explains an additional 2% of variance on average (median, p<1E-5) (**Figure 6C**).

#### Hidden states capture additional variance of neural encoding in orbitofronal cortex

As a second test that the states inferred from the model fit to choice behavior reflect meaningful computations, we investigated whether their dynamics correlate with changes in neural encoding. As a high-level test of this possibility, we apply the MoA-HMM to four animals that performed the two-step task while simultaneously having activity recorded in orbitofrontal cortex (OFC), data first published by ***Miller et al. (2022***). Using a single-state MoA model to estimate choice and outcome values, ***Miller et al***. found that OFC neurons primarily encoded the value of the most recently experienced outcome (or expected outcome value), used for learning about choice values on sub-sequent trials, during the ITI. Additionally, significant modulation to reward was observed in OFC neurons. Both reward and expected outcome value are relevant for learning between trials; therefore, our next analyses will focus on whether state modulates the encoding of these variables during the ITI.

**Figure 7A** looks first at OFC modulation to expected outcome value. For more direct comparison to results from ***Miller et al***., and since MBr is the largest contributor to behavior, expected outcome value is defined here as the value of the experienced outcome port predicted solely by the MBr agent prior to value updating following reward receipt. To look at a single unit’s response to expected outcome value, we binned trials by outcome value into terciles of outcome value. **Figure 7A** shows an example unit especially responsive to outcome value. Splitting the average firing rate by the inferred latent state reveals the highest modulation to outcome value during the second state (middle panel). To test the prevalence of this effect across all units, we measured the population CPDs of relevant behavioral variables in a Poisson regression model predicting the spike counts of each neuron computed separately for trials associated with each state. Specifically, our model included main effects of choice, outcome, transition (or interaction between choice and outcome), reward, next choice, expected outcome value, and next chosen value (see methods for additional detail). Looking at the CPD of expected outcome value split by state (**Figure 7B**) reveals that the trend from the example neuron is consistent across the population of OFC units, where state 2 has a significantly greater CPD than states 1 and 3. This is consistent with the idea that the second state generally has the highest weight on, and therefore the strongest influence by, the MBr agent.

**Figure 7.**
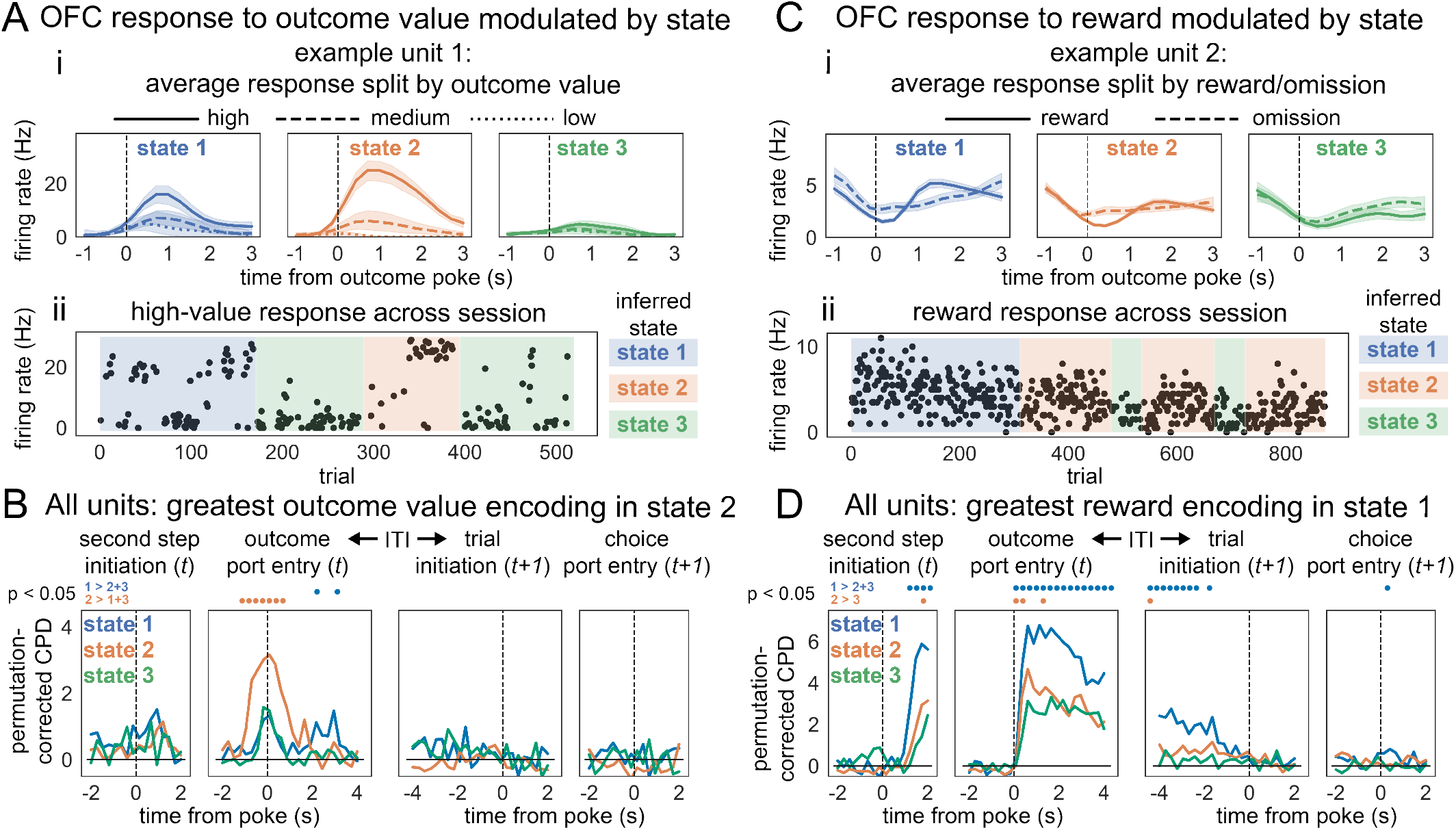
Orbitofrontal cortex (OFC) neural encoding modulated by state. **A** Modulation of model-based (MBr) outcome value encoding by state. (**i**) Peristimulus time histograms (PSTHs) of an example OFC unit time-locked to outcome for each state, where outcome time is measured as the time at which the rat enters the second-step outcome port. To differentiate between levels of expected outcome value, trials were split into terciles by value. Shaded areas are 95% confidence intervals around the mean. The greatest modulation to outcome value is seen in state 2. (**ii**) Average firing rate within the outcome window (defined from 1 second before until 3 seconds after outcome port entry) for each trial with a high expected outcome value throughout the session. Background shading corresponds to inferred state. Shifts in response magnitude align with shifts in inferred state. **B** Population CPD of expected outcome value throughout a trial, centered around the ITI, computed for each state. CPDs were baseline-corrected by subtracting out the mean CPD computed from circularly permuted datasets. Dots above each panel correspond to time points where a state’s CPD was significantly greater than the other states’ CPDs (blue: state 1 greater than states 2 and 3; orange: state 2 greater than states 1 and 3), i.e. when the CPD difference was greater than 95% of CPD differences computed from circularly permuted datasets. **C** Modulation of reward/omission encoding by state. (**i**) PSTHs of an example OFC unit time-locked to outcome for each state. To differentiate between responses to rewards and omissions, each PSTH is split by trials following omissions (dashed) and trials following rewards (solid). Shaded areas correspond to 95% confidence intervals around the mean. The first unit contains the highest response to reward in state 1 and the lowest in state 3. (**ii**) Average firing rate within the outcome window (defined from 1 second before and 3 seconds after outcome port entry) for each rewarded trial throughout the session. Background shading corresponds to inferred state. Shifts in response magnitude align with shifts in inferred state above a slow negative drift across the session. **D** The CPD of reward across all units throughout a trial, centered around the ITI, computed for each state, baseline corrected as in (B). Dots above panels similarly represent time points where a state’s CPD is significantly greater than the other states (blue: state 1 greater than states 2 and 3; orange: state 2 greater than state 3). **Figure 7—figure supplement 1**. Population CPD computed around the inter-trial interval (ITI) reveals significant encoding of state (gold) even after accounting for time (gray). CPDs were measured as the median CPD across all units, with errorbars corresponding to bootstrapped 95% confidence intervals. **Figure 7—figure supplement 2**. Example and summary 3-state MoA-HMM fits for rats with electrophysiological recordings in OFC.

**Figure 7C** looks next at OFC modulation to reward or omission. A second example unit (**i**) shows differentiable firing response to these events. Splitting response by state reveals that the contrast in firing rate between reward and omissions is highest during the first state (blue) and nearly absent during the third state (green). Similarly across the population using the same regression model as above, we see the greatest reward CPD during state 1 (**Figure 7D**). This progression may reflect a change in reward modulation coincident with motivation and thirst/satiety. The animal will be thirstiest at the beginning of a session, and therefore most motivated by reward; the modulation of neural encoding by reward, then, follows accordingly.

Additionally, the modulation of the reward effect can also be seen between states 2 and 3 — immediately after outcome, we see a small but significantly higher modulation to reward during state 2 than during state 3. Although state 2 generally occurred in the middle of the session for the behavioral rats, states 2 and 3 trade off more competitively in the model fits of the rats with physiological recordings (Figure 7 – figure supplement 2), making it less likely that the difference between these two states is simply due to further passage of time. Indeed, the example unit in Figure 7Bii is a session that ends in state 2, coinciding with an uptick in response to reward. This suggests that reward response magnitude may be further modulated by how model-based or how engaged the animal is in the task, as captured by the choice model, that may be dissociable from the simple passage of time.

We wished to formally investigate the state effects captured in Figure 7A-B are not simply due to the passage of time. To dissociate significant effects of these correlated factors, time and state, we constructed another Poisson regression model containing main effects of behavioral variables choice, outcome, transition, reward, next choice, expected outcome value, and next chosen value alongside state and time. Notably, the model also included interaction terms between each behavioral variable with state and time, as well as additional interactions with reward, as a more thorough representation of possible effects being encoded by the population. Each variable’s CPD, then, was computed by removing the main effect and all interaction terms containing that variable (see methods for a full specification of factors).

**Figure 7 – figure supplement 1** highlights the average CPD of state (gold) and time (gray) across all units. Similar to the CPDs computed for ITI rate, time explains a greater proportion of variance than state. However, we still observe a significant CPD for state, suggesting that state identity explains additional variance not captured by (even rather elaborate nonlinear effects of) time alone. Hidden states detected by the MoA-HMM, therefore, seem to provide useful boundaries to epoch electrophysiological data, opening a door for further analysis and contrast between identified behavioral states.

## Discussion

Different regions and subcircuits of the brain are thought to subserve different decision strategies — ways to learn about or evaluate candidate actions. Thus, deciding what to do may involve weighting information from multiple such systems, where the relative contribution of certain learning rules can vary based on internal motivations and/or external environmental factors. Dynamics of this sort are often thought to explain phenomena such as the formation of habits with over-training and a shift between exploratory and exploitative choice modes. However, MoA models traditionally used descriptively to measure the influence of separable learning rules, especially in RL, are unable actually to capture such dynamics, but instead characterize all choices as arising from an IID mixture of agents. In addition to failing to capture key phenomena like habit formation endogenously in the models, this limitation may mask interesting, lesser-used strategies and limit interpretation of underlying neural correlates. Another family of models, the GLM-HMM, attempts to account for time-dependent changes in behavior; however, by only modeling observable environmental factors, it cannot capture changes in terms of more abstract learning rules thought to be used by the brain.

Here we introduce the MoA-HMM, combining mixture modeling of abstract learning rules with temporal dynamics that can capture discrete changes in these mixtures throughout behavior. We apply this model to rats performing a multi-step, reward-learning task and examine the dynamic contribution of various reinforcement learning rules. We successfully capture shifts in strategy corresponding to differing weights on included RL rules, reflecting a within-session progression between exploration, exploitation, and reduced engagement in the task. Furthermore, these shifts in strategies significantly predict variables not used to fit the model; specifically, response time is fastest during the exploitation state, even after accounting for drifts due to time. These shifts additionally capture changes in neural encoding, notably showing the highest response to MB value during exploitation.

The two-step task has been historically used to measure the trade-off between MB and MF reward learning strategies as a proxy for the trade-off between goal-directed and habitual behavior. The rat version of the task used here, however, has not shown any significant involvement of MF reward learning in choice behavior. Accordingly, studies using this version of the task have primarily focused on its application to studying goal-directed behavior via MB planning (***Miller et al., 2017, 2022***). The dynamic shifts in strategy uncovered here, however, suggest that behavior still may reflect a trade-off between goal-directed (outcome-state focused) and habitual (choice-focused) strategies, just not via the traditional MB/MF distinction. Instead, if we identify habitual behavior by choice-level perseveration as proposed by ***Miller et al. (2019***), then our results show emergence of habitual control toward the end of sessions with a rise in MFc. Interestingly, the rise of MFc isn’t coupled with an opposing decay of MBr (i.e., goal-directed exploitation). Instead, we see increasing MFc accompanied by increasing (i.e., less negative) MBc, which likely reflects a decline in goal-directed exploration. This pattern further suggests that the goals vs. habits trade-off is not simply mapped to MBr/MFr as traditionally modeled. Instead, goal-directed contributions to behavior further subdivide into distinct MB exploratory and exploitative behaviors.

The nature of the MB/MF trade-off could be further explored by applying the MoA-HMM to other versions of the two-step task that do demonstrate significant contributions of MBr and MFr agents in the single-state MoA model (***Daw et al., 2011; Gillan et al., 2016; Hasz and Redish, 2018; Groman et al., 2019b; Akam et al., 2021***). This may uncover new properties of strategy switching when both MBr and MFr learning rules explain significant variance, especially in humans. One practical issue, and opportunity for future research, is that human datasets contain many fewer choices per subject than the rat dataset studied here. Thus to address these data, it would likely be necessary to develop a fully hierarchical MoA-HMM to pool data effectively over subjects.

Patterns of switching, in these and other cases, may also shed light on controversy around the interpretation of behaviors explained by mixtures of MBr and MFr agents in this task (***Akam et al., 2015; Russek et al., 2017; Feher da Silva and Hare, 2020***), e.g., whether they really reflect two separable contributions vs. some third hybrid, or an altogether different switching strategy. Indeed, it has been suggested that seemingly MB rat behavior during various rodent versions of the two-step task could instead reflect MF belief updates that switch between two latent states, one corresponding to each rewarded side. Such switching could be behaviorally indistinguishable from MBr in single-state MoAs for reasons similar to the example we show in Figure 3C (***Akam et al., 2015***). The MoA-HMM might instead be expected to capture this pattern via two strategies with opposing biases that track reward block flips. However, this is not what we observe in the animals. Even when aligning states 2 and 3 by signs in the bias, state dynamics do not follow reward block shifts (Figure 5 – figure supplement 1). While this does not necessarily rule out the interpretation of MF behavior put forward by ***Akam et al. (2015***), the emergent role of MBc describing changes in MB exploration throughout sessions provides additional evidence that the animals are MB in the sense of using transition information to inform decisions.

The emergence of putative exploratory and exploitative states presents a new opportunity for using this task to study the explore-exploit trade-off. First, the predominance of the MBc agent at the start of each session strongly suggests an interpretation for the function of the previously described “novelty preference” agent in directed exploration. In this sense, MBc (with a negative mixture coefficient) is analogous to including an “information bonus” for directed exploration, similar to models used to describe exploratory behavior previously (***Wilson et al., 2014, 2021***). Such a bonus, in turn, captures the qualitative features of more elaborate, exact decision-theoretic analyses of the value of exploration (***Gittins, 1979; Dayan and Daw, 2008; Averbeck, 2015; Costa et al., 2019; Hogeveen et al., 2022***). In a different way, our results echo another recent study (***Ebitz et al., 2018***) which argued that primate behavior and neural responses were well-described by a (more purpose-built) HMM that switched between discrete exploratory and exploitative choice regimes.

Importantly, unlike single-step “bandit” tasks used in these and other studies of exploration, the two-step task is sequential. This means that effective exploration requires not simply choosing actions about which one is directly uncertain (as with MFc in the present task) but instead choosing actions that *lead to future states* (here, outcome ports) whose rewards are uncertain; this is precisely what distinguishes MBc as model-based. In this sense, the two-step task allows for — and our result of negative MBc but no negative MFc shows to our knowledge the first evidence in animal or human behavior for — what is known in RL as “deep exploration” (***Wilson et al., 2020***), i.e., behavior targeted at reaching future states for the purposes of exploring them.

Following the GLM-HMM, HMMs have been utilized recently in reinforcement learning tasks to estimate strategy switching (***Le et al., 2023; Li et al., 2024***). However, these approaches are notably different than the MoA-HMM. Specifically, the blockHMM designed by ***Le et al***. captures differences in choice-switching behavior during a reward reversal task, and is therefore limited in application against tasks that do not contain reward reversals. Furthermore, the MoA-HMM uses different cognitive models to directly infer strategy switching, where ***Le et al***. decodes underlying algorithms post hoc to states identified by the blockHMM, where decoding is limited to mixtures of only two algorithms (model-free and inference-based learning) relevant to reward reversal tasks. Conversely, dynamic noise estimation by ***Li et al***. extends more broadly to various tasks and cognitive models. However, this method only assumes two behavioral states, engaged or disengaged, in order to improve modeling of the “engaged” state by separating out noise from the “disengaged” state. On the other hand, the MoA-HMM can extend to an arbitrary number of states and does not assume the existence of a “disengaged” state. Furthermore, instead of attributing strategy switching to “noisy” deviations from a single cognitive model, the MoA-HMM can instead identify competing cognitive models that may underlie strategy switches.

As a descriptive model, the MoA-HMM assumes discrete rather than continuous strategy shifts. It is possible that our results reflect a discrete approximation to a more continuous process. Our goal is to provide a quantitative description of significant changes to behavior, which may still offer a useful window into continuously changing strategies. A way to more explicitly assess discrete vs. continuous dynamics is to compare the fit of the MoA-HMM to that of a behavioral model with continuously drifting latent states, such as Psytrack (***Roy et al., 2021; Ashwood et al., 2020***), which could be adapted (much like GLM-HMM was extended to MoA-HMM) to infer continuous shifts in learning rule mixture weights.

The appearance of switching dynamics might also arise if there is a single underlying agent whose continuous dynamics (e.g. algebraic value learning rules) do not match the MoA agents.

In this case, HMM states might capture residual variance due to this mismatch. In this case also, the current model would still be useful descriptively (e.g., to help design better underlying agents that endogenize the dynamics described by the HMM). Anecdotally, we were not able to capture switching in the present dataset by exploring agent variants, and at least some of the dynamics we observe (e.g. motivational shifts) seem unlikely to reflect learning per se. Nevertheless, an approach to address this possibility in future work would be to pair the MoA-HMM with more flexible agent models that seek to infer the continuous learning rules being used (***Eckstein and Collins, 2021; Miller et al., 2023; Ji-An et al., 2023; Correa et al., 2024***).

Finally, there are many promising applications of the MoA-HMM that have yet to be explored. For example, while the MoA-HMM itself is a descriptive model of behavior and is not explicitly modeling an underlying arbitration of controllers in the brain, the resulting behavioral states may be indicative of underlying neural processes and help identify times when different neural controllers are prevailing. Therefore, to test whether separable circuits support different strategies, temporally-specific perturbations targeting these circuits could test for isolated effects on the agent or agents describing said strategy. Additionally, finding a significant relationship between MoA-HMM states with behavioral response times and neural encoding alone cannot causally link the observed effects with inferred states. However, it does suggest that the inclusion of these additional metrics may provide a richer description of changing strategy that modeling choice alone cannot capture. Indeed, models combining both behavioral and neural data have found success in capturing behavioral and neural patterns that could not be captured by one modality alone (***Shahar et al., 2019; DePasquale et al., 2022; Luo et al., 2023***). Furthermore, while the MoA-HMM has been applied here in the context of RL, the framework can be applied more generally to arbitrary learning rules and tasks, and it would apply naturally to rule-switching or task-switching settings.

## Methods and Materials

### Subjects and Behavior

The subjects used in this study were drawn from two published data sets. For behavioral analyses, we used 20 rats first published by ***Miller et al. (2017***); for analyses paired with electrophysiological recordings, we used 4 rats first published by ***Miller et al. (2022***). All subjects from both sources underwent the same training procedure. Briefly, rats were water deprived and trained daily on the two-step task to gain their daily allotment of water (Figure 1A). Behavioral training took place in custom behavioral chambers (Figure 1A, bottom right), and each behavioral session lasted 2-3 hours during which the animal completed hundreds of trials. Before a rat would train specifically in the two-step task, they would undergo multiple shaping phases to 1) acclimate them to receiving water reward from successfully poking into the back-lit outcome port, 2) train them on the transition structure of the task via fully-instructed trials (where transition structure is fixed within rat and counterbalanced across rats), and 3) introduce free choice trials and reward reversals, where reversals only occurred after a performance threshold was reached. Within this stage, the reward reversal probabilities were walked down from 100%/0% to the final 80%/20% probability set. Once the rat was consistently triggering multiple reversals per session, rats were advanced to the full version of the task where reward reversals could trigger regardless of the performance of the animal. Notably, some rats still have a small percentage of forced-choice trials during the full task (up to 20%) to reduce strong side biases. For further details on the training pipeline, see ***Miller et al. (2017***).

### Mixture-of-Agents Model

The mixture-of-agents model used here was taken and extended from ***Miller et al. (2017***). This model describes behavior as arising from a mixture of multiple agents, *A* ∈ {agents}, each with their own method to calculate an choice value *Q*_*A*_(*y*) for each choice *y*. The MoA uses a weighted sum of each agents’ values to calculate choice likelihood for each *y* according to the softmax function,

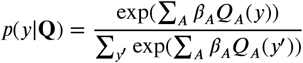

where *β*_*A*_ is the weighting parameter (or inverse temperature) on agent *A*. Importantly, forced-choice trials are *included* in updated choice values *Q*_*A*_(*y*), since the animal can still observe and learn from rewards during these trials; however, forced-choice trials are *excluded* when calculating total choice likelihood to infer the weighting parameters (fitting procedure outlined in Mixture-of-Agents Hidden Markov Model section).

#### Model Agents

We use five agents in the present manuscript. Four of these agents are separable reinforcement learning rules to capture model-based and model-free influences of of reward and choice, derived from learning rules introduced by ***Ito and Doya*** (***2009***).

### Model-Based Reward Learning

The model-based reward (MBr) agent captures planning as the interaction of reward and transition on choice; however, the task transition probabilities, which are assumed to be known by the animals, have been simplified to definite transitions, such that common transition choices are updated directly from reward experienced at the second step. After each trial *t*, the choice values for each *y, Q*_MBr_(*y*), are updated according to

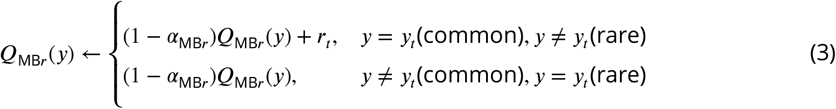

where *α*_**MBr**_ is a decay rate/learning rate on the influence of previous choice value, constrained between [0, 1], and *r*_*t*_ is an integer representing the reward experienced on trial *t*, taking a value of [1, −1] for rewards and omissions, respectively. With the transition requirement on *y*, this learning rule will apply the reward value to selected choice (*y* = *y*_*t*_) and decay the non-selection choice (*y* ≠ *y*_*t*_) following common transitions, or apply the reward value to the non-selected choice (*y* ≠ *y*_*t*_) and decay the selected choice (*y* = *y*_*t*_) following rare transitions. Notably, *α*_MBr_ is not applied as a learning rate on reward *r*_*t*_ to reduce interaction with the weighting parameter *β*_MBr_ during model fitting, which is similarly applied in the following RL agents.

### Model-Based Choice Learning

Model-based choice (MBc, also model-based perseveration or outcome perseveration) captures the weighted influence of past transitions on choice, describing a “common-stay, rare-switch” pattern of choices. After each trial *t*, choice values *Q*_MBc_ are updated as

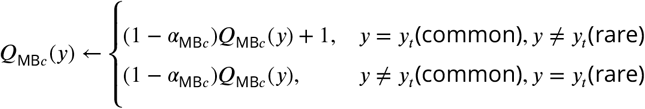

where *α*_MBc_ is a decay rate, similarly constrained between [0, 1], which is unique from the decay rate on the previous agent, MBr. With a similar transition requirement on choice, this agent will always add 1 to selected choices (*y* = *y*_*t*_) following common transitions or non-selected choices (*y* ≠ *y*_*t*_) following rare transitions, regardless of reward value. This amounts to reinforcing experienced outcomes by reinforcing the common transition to that outcome.

### Model-Free Reward Learning

Model-free reward learning (MFr) is equivalent to a temporal difference learning rule, specifically TD(1), where the choice values are updated directly from experienced reward at outcome time. Namely, choice values *Q*_MBr_ are updated according to

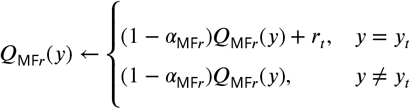

where *α*_MFr_ is a decay rate constrained between [0, 1], distinct from both model-based decay rates. With transition no longer a contributing factor, this agent will always apply the experienced reward value *r*_*t*_ to the selected choice (*y* = *y*_*t*_) and always decay the non-selected choice (*y* ≠ *y*_*t*_).

### Model-Free Choice Learning

The model-free choice agent (MFc, also model-free perseveration or choice perseveration) captures the animal’s tendency to repeat choices on subsequent trials. Specifically, choice values *Q*_MFc_ are updated as

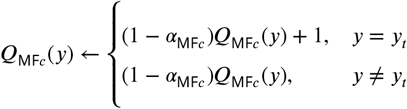

where *α*_MFr_ is a decay rate, also constrained between [0, 1], unique from the decay rate on MFr. This agent will always reinforce selected choices (*y* = *y*_*t*_) regardless of outcome or reward.

### Bias

The bias (or intercept) agent captures the animal’s overall tendency towards left or right choices, which has a set value *Q*_bias_ of

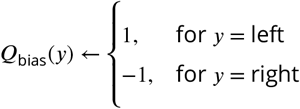

that stays constant across trials.

#### Predecessor agents from ***Miller et al. (2017***)

The agents used in the original MoA model used for the rat two step task are described in ***Miller et al. (2017***). In Figure 4–figure supplement 1, we show that our extended learning rules show an improvement in model likelihood over these original agents. We include a brief description of these four agents below, and additionally a fifth model-free agent used in the original manuscript (but not in the reduced model) that we included for fairness in model comparison.

### Model-Based Temporal Difference Learning

Computationally similar to MBr, this agent is equated to planning, which uses knowledge of the transition structure between choices and outcomes to update choice values. Specifically, the value of choice *y, Q*_MB_(*y*), is updated as the weighted sum:

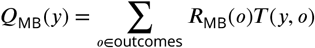

where *R*_MB_(*o*) is the value of the outcome *o*, and *T* (*y, o*) is the transition probability from choice *y* to outcome *o*, which is assumed to be known. The value of outcome *o* is updated using a TD learning rule as follows:

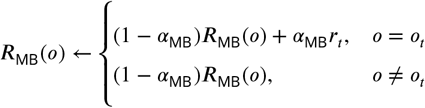

*r*_*t*_ is a constant indicating reward observed on trial *t* which takes the value [−1, 1] for omission and rewarded trials, respectively, and *α*_MB_ is the learning rate which takes a value between 0 and 1. Reward value is used to learn about the experienced outcome *o* = *o*_*t*_, and the non-experienced outcome *o* ≠ *o*_*t*_ is decayed.

### Novelty Preference

Using the same terminology and definition as ***Miller et al. (2017***), this agent captures the tendency to stay following a rare transition or switch following a common transition. The selected choice value *Q*_NP_(*y*_*t*_) and non-selected choice value *Q*_NP_(*y* ≠ *y*_*t*_) are updated as:

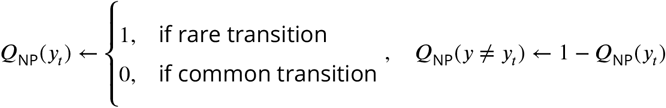

This agent is similar to MBc,except that it is defined in the opposite direction (i.e. “rare-stay/common-switch”) and has no learning rate to capture trial history effects.

### Model-free Temporal Difference Learning

This agent is nearly identical to MFr, where choice values are updates as:

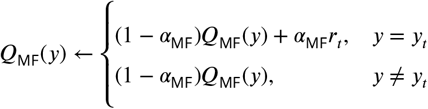

Unlike MFr, however, the learning rate *α*_MF_ is applied to reward value *r*_*t*_ for consistency in definition with ***Miller et al. (2017***).

### Perseveration

The perseveration agent captures the animal’s tendency to repeat choices on subsequent trials. its value is update according to

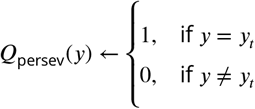

This agent is similar to MFc, except that is has no learning rate to capture trial-history effects.

### Mixture-of-Agents Hidden Markov Model

The introduction of a hidden Markov model allows for the weighted influence of each agent to change discretely over time. Specifically, it assumes that behavior is described by multiple hidden states, each with their own set of agent weights ***β***^*z*^, of which one set will win out on any given trial to drive choices. Our MoA choice likelihood is updated, then, to be dependent on latent state such that:

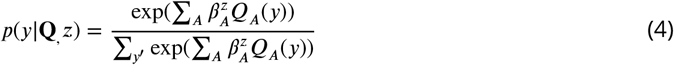

where the weighted influence of each agent *β*^*z*^ is additionally dependent on latent state. We assume that all latent states use a shared value *Q*_*A*_(*y*) and learning rate s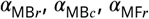 and 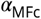.

Two additional parameters are introduced from the HMM to describe the temporal dynamics of the hidden states: the initial state probability *π* and the hidden state transition matrix **P**, making the full set of model parameters 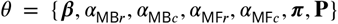 where ***β*** is the full (nAgents x nStates) weight matrix. Details on the inference procedure for each of these parameters follow.

Initial state probability *π* is typically defined as the hidden state probability distribution on the first trial, *p*(*z*_1_). However, with multiple sessions with only a single session a day, we treat the first trial within a session as an “initial” trial. Initial state probability *π* is estimated, then, from all trials that initiate a session. As stated previously, forced trials are excluded from choice likelihood estimation. For sessions that start with a forced trial, the first free choice trial is considered the initial trial of that session. We can break up the set of all trials *T* into subsets of trials belonging to each session *s* ∈ *S*, for all sessions *S*, such that 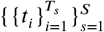, where *i* = 1 is the first trial and *i* = *T*_s_ is the last trial in session *s*; therefore, the subset of all initial trials is denoted 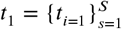. Elements of *π* are defined as the average probability

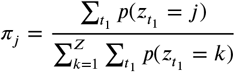

The transition matrix is estimated from all trials excluding those that initiate a session, denoted as 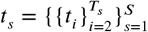. Elements of **P** are 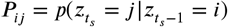

We fit the MoA-HMM using expectation maximization (EM), which finds the posterior distribution of the hidden state variables, *p*(*z*|*D, θ*^*old*^), in the expectation (E) step to evaluate and optimize the expected data log-likelihood, *𝓁𝓁*(*θ, θ*^*old*^) = Σ_*z*_ *p*(*z* | *D, θ*^*old*^) ln *p*(*D, z* | *θ*), in the maximization (M) step. Implementation closely follows ***Bishop (2006)***, which we briefly describe to highlight differences.

We introduce the following notation: *γ*_*tj*_ = *p*(*z*_*t*_ = *j*|*D, θ*^*old*^), the marginal posterior probability of hidden state *z*_*t*_ = *j* at trial *t*; and *ξ*_*tij*_ = *p*(*z*_*t*−1_ = *i, z*_*t*_ = *j*|*D, θ*^*old*^), the joint posterior distribution of two successive latent variables, *z*_*t*−1_ = *i* and *z*_*t*_ = *j* at trial *t*. With these definitions, our expected data log-likelihood becomes:

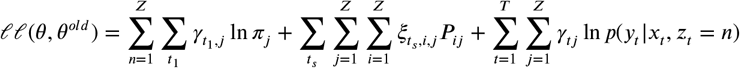

where *T* is the total number of trials and *Z* the total number of hidden states. The E step will evaluate *γ* and *ξ* using existing parameter values *θ*; Optimization of *θ* during the M step is then done by treating *γ* and *ξ* as constants.

### E Step

*γ* and *ξ* are estimated using the forward-backward algorithm as described by ***Bishop*** (***2006***). Briefly, the forward pass is calculated recursively within sessions from *t* = 2 to *t* = *T*_*s*_ as:

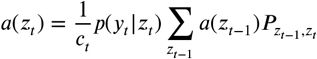

Where *c*_*t*_ is a normalization factor such that 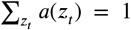, which also gives the marginal choice likelihood 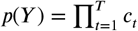. Notably in our case, *a* is initialized at the beginning of every session as:

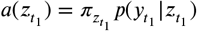

The backward pass is calculated backward-recursively within sessions from *t* = *T*_*s*_ − 1 to *t* = 1 as:

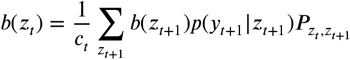

Where *c*_*t*_ is the same normalization factor calculated from the forward pass. Similarly, *b* is “initialized” at the end of each session as 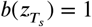

*γ* is calculated as:

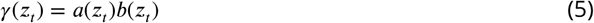

Importantly, the value of *γ*(*z*_*t*_) gives us the expected likelihood of state *z* on trial *t*. Finally, *ξ* is calculated as:

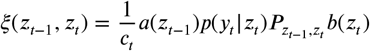

### M Step

Since *π* and **P** do not depend on hidden state *z*, they can be maximized using Lagrange multipliers:

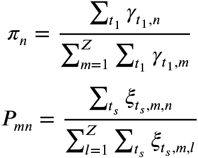

A notable difference is the calculation of *π* includes all initial trials for each session as defined before, while **P** excludes these trials.

Dependence on agent parameters and agent weights 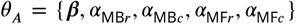 are limited to a single term in our data log-likelihood:

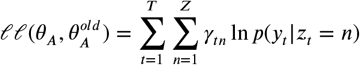

the negative of which can be optimized using gradient descent. Here we use the L-BFGS algorithm implemented by julia’s Optim package, making use of automatic differentiation to supply the gradient.

#### Fitting Procedure

Parameter values were randomly initialized using the following distributions: the agent weights ***β*** from a normal distribution *N*(0, 0.1); transition matrix rows from a Dirichlet distribution, where shaping parameters were taken as the corresponding row from a *Z* x *Z* matrix of 1s with 5s along the diagonal; the initial state probability from a Dirichlet distribution with all shaping parameters = 1; the learning rates from Beta distribution with *α* = *β* = 5. In addition, a loose prior on ***β***, denoted *p*(***β***), was included to penalize large magnitudes; explicitly, the prior was a normal distribution with mean 0 and standard deviation 10. Including a parameter prior updates the data log-likelihood used in the M Step to

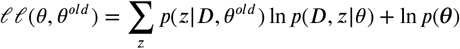

where *p*(*θ*) = *p*(***β***) = Normal(0, 10) with only a prior on ***β***. EM was terminated if the difference in the maximum a posteriori (MAP) likelihood, *p*(*θ*| *Y*) ∝ *p*(*Y*) *p*(*θ*), fell below 1 × 10^−5^, where *p*(*Y*) is the marginal choice likelihood computed from the forward pass during the E step, and *p*(*θ*) is the parameter prior, which at this point *p*(*θ*) = *p*(***β***) = Normal(0, 10). Alternatively, if the difference in all parameter estimates on subsequent iterations fell below 1 × 10^−5^, or if the number of iterations exceeded 300, EM would be terminated.

#### Estimation of Population Prior

Hierarchical modeling gives a better estimate of how model parameters can vary within a population by additionally inferring the population distribution over which individuals are likely drawn (***Daw, 2011***). This type of modeling, however, is notoriously difficult in HMMs; therefore, as a compromise, we adopt a “one-step” hierarchical model, where we estimate population parameters from “unconstrained” fits on the data, which are then used as a prior to regularize the final model fits. This approach is motivated by analogy to the first step of EM fit in a hierarchical model, in which population-and subject-level parameters are iteratively re-estimated in terms of one another until convergence (***Daw, 2011; Huys et al., 2012***). It is important to emphasize, since we aren’t inferring the population distributions directly, that we only estimate the population prior a single time on the “unconstrained” fits as follows.

The best model was selected from at least 20 fits with random initializations for each rat using only a loose prior on the agent weights (nearly unconstrained). The best fits were sorted according to their initial state probability for the first state, and by *β*_MBr_ magnitude for the remaining two states, and used to estimate prior distributions on each of the model parameters. The model was refit using the updated prior:

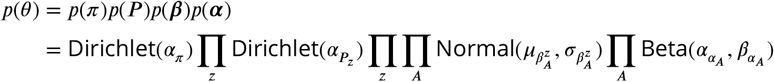

The prior on *β* was a Normal distribution for each state, with mean 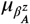 and standard deviation 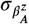 for each agent *A* and state *z* approximated from the sorted population parameters.

The prior on *π* and each row of **P** are Dirichlet distributions, where the vector elements of *α*_π_and 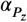 are estimated from population parameters using the method of moments:

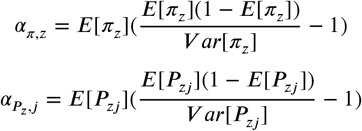

where *j* can be any element of row *P*_*z*_. To ensure all shaping parameters were at least 1, 1 was added to each index of *α* if any index was less than 1, effectively smoothing the prior. To account for the prior, closed form updates to *π* and **P** were updated as:

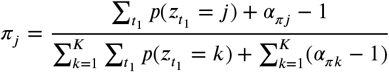

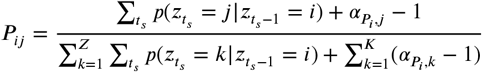

The prior on the learning rates was taken as a Beta distribution, with shaping parameters 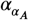 and 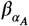 estimated for each RL agent also using the method of moments:

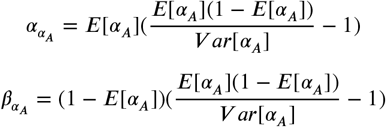

Similarly as before, if either 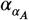 or 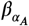 were estimated as a value less than 1, 1 was added to both 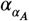 and 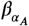.To reduce the number of iterations and for further bias towards the prior, making up for the fact that the hierarchical estimates of the prior aren’t being inferred alongside each model, the model was refit for each animal with a single initialization at the population mean.

### Data simulation and parameter recovery

To test the model’s ability to recover inferred parameter sets, we generated two-step behavioral data from a given MoA-HMM. Given a number of trials evenly split between a number of sessions, reward probabilities were first generated for left and right outcomes using task statistics: on the first trial, a either left or right was randomly chosen and assigned an 80% reward probability, and the opposite side assigned a 20% reward probability. A trial flip counter, initialized to 0 at the start of every session, was used to track the number of trials following a reward probability flip. After the trial flip counter reached 10 trials, a random number was drawn on each trial to determine whether the reward probabilities would flip sides, with flips occurring at a 5% probability. The trial flip counter was reset to 0 following a successful flip, and reward probabilities were generated in this manner throughout the set of simulated trials and sessions.

After generating reward probabilities, choices were generated from the probability distribution given by equation 4 using a given MoA-HMM parameter set. Similarly to model inference, choice values were initialized uniformly at the start of every session. After a choice was drawn, a common or rare transition was randomly chosen based on an 80%/20% transition probability to determine the outcome side, and reward was drawn from the predetermined reward probabilities for the given outcome. The selected choice, outcome, and reward are used to update choice values, and this process is repeated for the specified number of trials and sessions.

The MoA-HMM was then fit to the simulated data set using the model fitting procedure outlined above, finding the best model fit out of at least 5 random initializations. In order to match recovered HMM states to the simulated HMM states, we computed the expected state likelihood *γ* (equation 5) for each state across the simulated data set for both the simulated model, *γ*_sim_, and the recovered model, *γ*_fit_, and we paired states based on the maximum correlation between *γ*_sim_ and *γ*_fit_. If multiple states in *γ*_fit_ were maximally correlated to the same state in *γ*_sim_, the state with the highest correlation won out, and the losing state was paired with its next highest correlated state. This process was repeated until all states have unique pairs.

Parameter recovery was performed separately on two sets of models: 1) models with randomly drawn parameter sets, and 2) models inferred from the rats’ behavior. For the first set, model parameters were randomly drawn from user-defined distributions. Specifically, agent weights *β* for all agents were drawn from a Normal distribution with *μ*_*β*_ = 0 and *σ*_*β*_ = 2. Learning rates *α* for relevant agents were drawn from a Beta distribution with shaping parameters *α*_*α*_ = 5 and *β*_*α*_ = 5. Initial state probabilities *π* were drawn from a Dirichlet distribution with shaping parameter *α*_*π*_ = [1, 1, 1], and each row of the state transition matrix *P* was also drawn from a Dirichlet distribution using the corresponding row of a shaping parameter matrix with 1 on all off-diagonals and 10 along the diagonal. 200 models were initialized under this framework, and data for each model was generated for 5000 trials evenly split between 20 sessions. Each model was recovered with no priors on any of the parameters except for a loose prior on the agent weights, Normal(0, 10).

Due to the interaction between different model parameters (e.g. a small *β* weight will affect the recoverability of the agent’s learning rate *α*), a number of “failures” can be seen in recovery of the first model group (Figure 3 – figure supplement 1). Instead of finding restrictions on the randomization of model parameters to eliminate failures, we performed parameter recovery on a second group of models defined by those fit to real data (post-prior) to ensure confidence in our ability to recover these models specifically (Figure 3 – figure supplement 2). Data generated by these models were set to match the trial and session statistics of the rat’s behavioral data from which the model was inferred as a further check on the recoverability from the size of the data set. 5 independent data sets were generated from each of the 20 inferred models, giving a total of 100 recovered sets. Similar to the fitting procedure on each rat, the prior distributions computed from the pre-prior model fits were used when fitting the model to each simulated data set.

### Cross-validation and model comparisons

Cross-validation (CV) was used for all procedures comparing models with different parameter sets. Either 3- or 6-fold cross validation was used, where entire sessions were held out together to preserve within-session temporal structure. Sessions for each fold were selected in stride, such that sessions were evenly spaced throughout the data set (i.e. every third session belonged to the same set in 3-fold CV). The CV likelihood, then, was computed by fitting the model to all data excluding the held-out set, and then running the forward pass of the E-step to compute the marginal choice log-likelihood on the held-out set, ln *p*(*Y*_*set*_). After being computed for each fold, each fold’s log-likelihood was added and normalized by the number of trials, and exponentiated back into likelihood space, giving the model-fit likelihood:

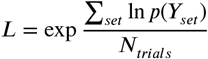

### GLM-HMM parameter description

The GLM used in the GLM-HMM version of the model used here was taken from ***Miller et al. (2017***). It consists of trial-history predictors that indicate what trial type was experienced at multiple delays. Specifically, there is a common-reward predictor that takes a value of 1 if the trial was a left-choice rewarded trial with a common transition, −1 for a right-choice rewarded trial with a common transition, and 0 otherwise. This pattern follows for the remaining three trial types: common-omission, rare-reward, and rare-omission. Each of these four predictors are measured at variable delays, i.e. one trial back up to five trials back; including a bias term, this gives a total of 21 predictors for each state. This model was fit similarly to the MoA-HMM, with each predictor treated as an “agent” with no agent hyperparameters to infer.

### Multilevel linear regression of response times

Multilevel linear regression was used to understand the influence of reward, state, and time on ITI duration. ITI duration was first pseudo-linearized by transforming it into a rate, or 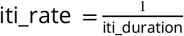. The relationship between these variables can be summarized by the following formula:

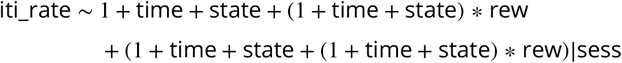

“Time” consists of three terms, time = *t* + *t*^2^ + *t*^3^, where *t* is the initiation time of the trial, min-maxed normalized across the session such that 0 corresponds to the beginning of the session and 1 corresponds to the end of the session. Time is described by a third-degree polynomial in this way to capture the possibility of asymmetric, non-linear effects of time throughout the session.

“State” consists of two terms, state = *z*_1_ + *z*_2_, where *z*_1_ and *z*_2_ are non-overlapping boolean vectors. *z*_1_ = 1 indicates state 1 is the most-likely inferred state, and *z*_2_ = 1 indicates state 2 is the most-likely inferred state. A third boolean vector for state 3 is implicit with the intercept and is therefore excluded. In other words, “state” is a one-hot matrix corresponding to hidden state, where the column corresponding to the third state is dropped.

“Rew” (short for “reward”) is a boolean vector, where 1 corresponds to trials following reward and 0 corresponds to trials following omissions. The interaction between each time and state term with reward was also included in the model

The regression was fit individually for each rat, with random effects of session (“sess”) on each of the included terms to account for variability across sessions. Regressions were fit using the MixedModels package in julia.

### Coefficient of partial determination

The coefficient of partial determination (CPD) measures the variance explained by a regressor (*x*) that cannot be accounted for by other potentially correlated regressors. The CPD is defined as the proportional change in the residual sum of squares (RSS) to the held-out model (RSS_−*x*_) from the full model (RSS_full_).

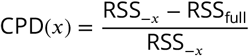

The CPD can take a negative value if the RSS of the held-out model is less than the RSS of the full model.

### Non-parametric statistics

Since the distribution of behavioral and model metrics were non-normal and calculated at the level of rats, we performed non-parametric statistics to determine significant differences between groups. Specifically, we used a Wilcoxon signed-rank test on matched samples. These tests were performed using the SignedRankTest function in the HypothesisTests package in julia.

### Electrophysiological recordings and PSTH computation

Electrophysiological data were recorded from OFC in four rats, described by ***Miller et al. (2022***). Briefly, 32-channel microwire arrays (Tucker-David Technologies) were surgically implanted to unilaterally target OFC (coordinates: 3.1-4.2 mm anterior, 2.4-4.2 mm lateral relative to bregma; lowered 5.2 mm from bregma or about 4.2 mm below the brain surface). After recovery, recording sessions were acquired in a behavioral chamber outfitted with a Neuralynx recording system. Spiking data were detected from 600-6000Hz bandpass filtered voltage traces using a threshold of 30*μ*V and manually clustered.

Peristimulus time histograms (PSTHs) were computed for examples units to show changes in firing rate at outcome time due to differing task variables relative to each inferred state. Firing rates on each trial were computed in 250*ms* bins spanning 1 second before and 3 seconds after the time of outcome port entry, then smoothed using a Gaussian filter with kernel standard deviation of 1. In both cases, trials were initially split into three groups corresponding to the inferred state, measured as the state with the highest likelihood given by equation 5. For the first example, trials were further split by outcome value inferred by the MBr agent. The outcome value inferred by the MBr agent corresponds to the value described by equation 3 relative to the experienced outcome *before* the reward update has occurred. Specifically, on common trials, the outcome value of trial *t* is equivalent to the chosen value of trial *t, Q*_MBr_(*y* = *y*_*t*_), before the reward update; on rare trials, the outcome value of trial *t* is equivalent to the non-chosen value on trial *t, Q*_MBr_(*y* ≠ *y*_*t*_), before the reward update. Trials were split into three groups defined by low (−2 to −0.5), medium (−0.5 to 0.5), and high (0.5 to 2) outcome values, giving 9 total trial groups in the first example. For the second example, trials were further split into two groups corresponding to rewarded and omission trials, giving a total of 6 trial groups. PSTHs were computed as the mean firing rate within each trial group, and error bars computed as 95% confidence intervals around the mean.

### Poisson regression of neural activity and population CPD

Two Poisson regression models were used to predict spiking activity and calculate the CPD of constituent predictors. A simpler model was fit when calculating the CPD separately for each state only including main effects of each predictor and a single interaction between choice and outcome for transition, which can by summarized by the formula:

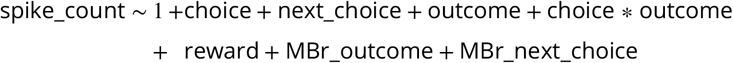

Spike counts were measured in 250*ms* time bins centered around task events: −2*s* before to 2*s* after second step initiation (16 bins), −2*s* before to 4*s* after the outcome port entry (24 bins), −4*s* before to 2*s* after the next trial initiation (24 bins), and −2*s* before to 2*s* after the next trial’s choice port entry. This regression was fit separately for each unit in each time bin, and the population CPD for each time bin *i* was computed as the aggregated CPD across all units *u* within that bin:

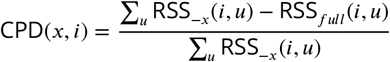

For this first model, to ensure that effects from the included predictors are not due to trial-by-trial co-fluctuations in neural activity, trial identities were circularly permuted and the CPD recomputed for each permuted session. A CPD (for a particular predictor in a particular state in a particular time bin) was considered significant if that CPD computed using the true dataset was greater than 95% of corresponding CPDs (same predictor, same state, same time bin) computed using these permuted sessions. For display, we subtract the average permuted session CPD from the true CPD in order to allow meaningful comparison to 0.

To test whether neural coding of a particular predictor in a particular time bin significantly differed according to HMM state, we used a similar test. For each CPD that was significant according to the above test, we computed the difference between that CPD and the CPD for the same predictor and time bin in the other HMM states. We compare this difference to the corresponding differences in the circularly permuted sessions (same predictor, time bin, and pair of HMM states). We consider this difference to be significant if the difference in the true dataset is greater than 95% of the CPD differences computed from the permuted sessions.

The second model, used to measure the separable effects of state and time, adds complexity by including not only main effects of time and state, but also interactions of time and state with every other predictor. Additional interactions of reward with outcome and next choice were also included.

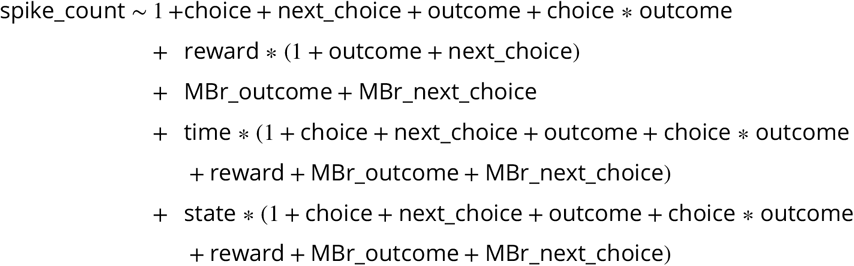

In the same way as the previously described model used to predict behavioral response times, the “time” predictor consists of three terms, time = *t* + *t*^3^ + *t*^3^, and the “state” predictor consists of two terms, state = *z*_1_ + *z*_2_. Therefore, all interactions listed above are repeated for each term in the “time” and “state” predictors. In order to calculate the CPD of each predictor, the main effect and all interaction terms were held out when fitting the reduced model. Explicitly for each predictor,

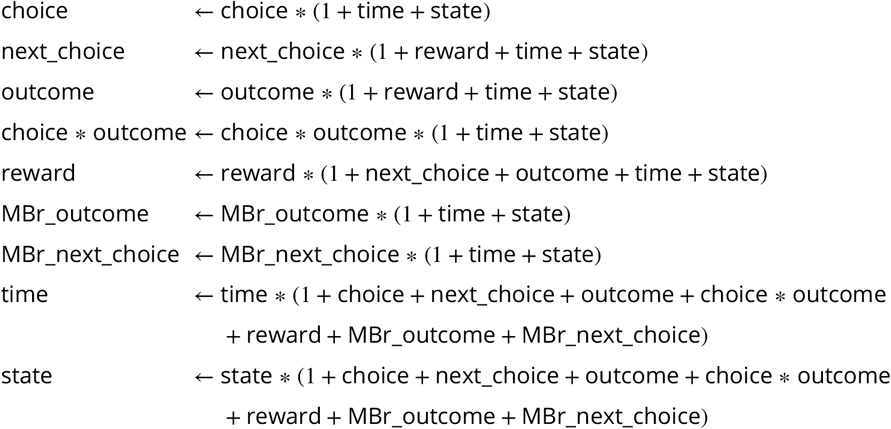

Since the effects of time and state *are* dependent on trial-by-trial co-fluctuations, permuting the trial identities does not provide a meaningful control; the inclusion of these interactions, therefore, is intended to bolster the description of possible time and state effects. Instead of reporting the aggregated population CPD of state and time, the median CPD across all units and 95% confidence intervals around the median were computed to measure significant variance explained by time and state. All Poisson regression models were fit using Scikit-Learn in python.

## Code availability

The MoA-HMM and figure code is available at https://github.com/Brody-Lab/MixtureAgentsModels

## Acknowledgments

We would like to thank Iris Stone and Jonathan Pillow for helpful discussions about GLM-HMMs, and Kim Stachenfeld for feedback on the manuscript. This material is based upon work supported by the National Science Foundation Graduate Research Fellowship Program under Grant No. DGE-1656466, a Research Innovation Award from Princeton Neuroscience Institute, and NIMH R01MH121093.

---

**Figure 3—figure supplement 1.**
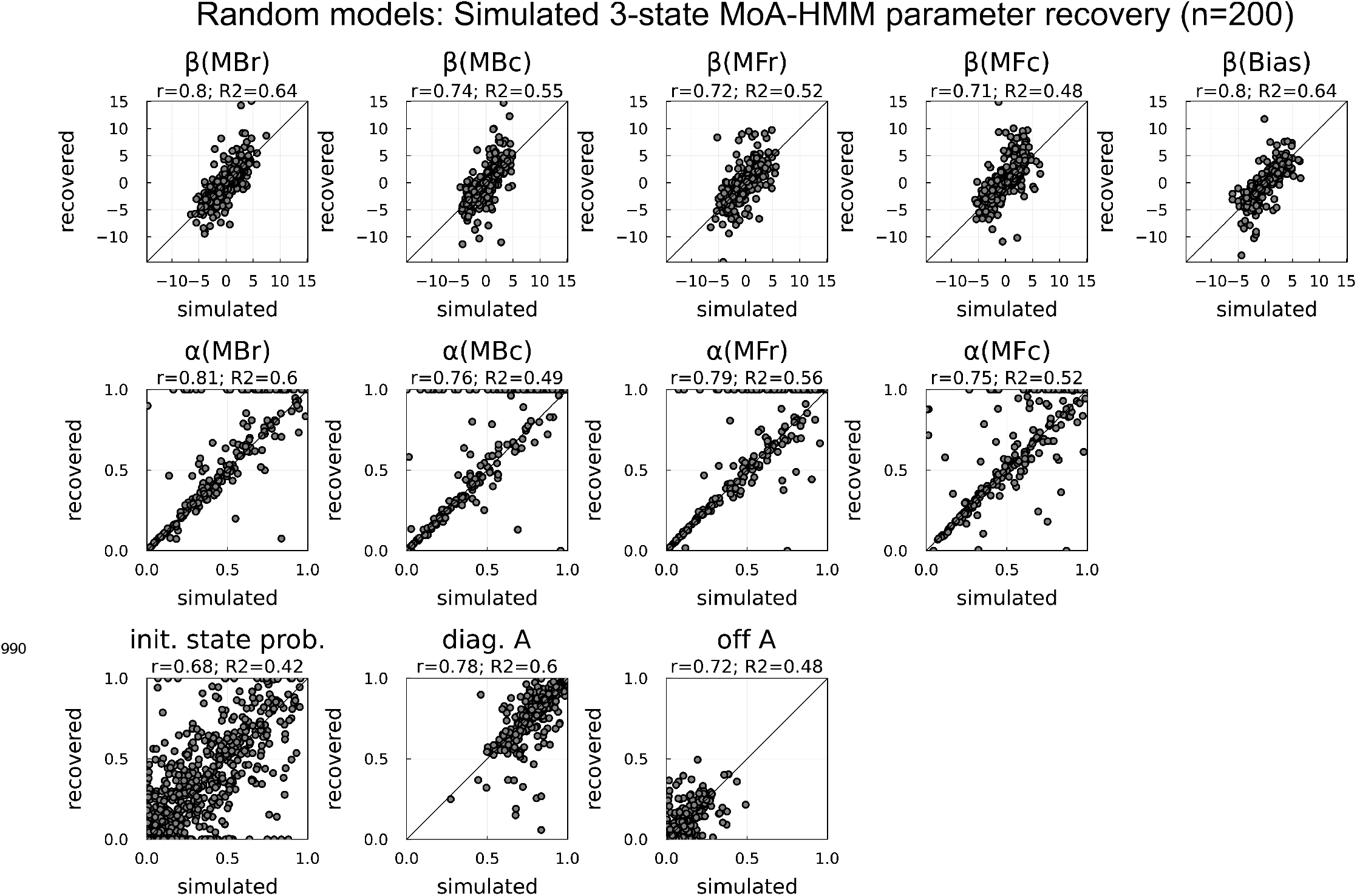
Simulated 3-state MoA-HMM parameter recovery for the five agents used in behavioral fits: model-based reward, model-based choice, model-free reward, model-free choice, and bias. Simulated model parameters were randomly drawn (see methods). Each simulation contained 5000 trials evenly split between 20 sessions. Parameters in each panel are pooled across states. “r” is the Pearson’s correlation coefficient between the simulated and recovered parameters, and “R2” is the coefficient of determination, *R*^2^, calculating how well the simulated parameters predict the recovered parameters. Due to the interaction between different model parameters (e.g. a small *β* weight will affect the recoverability of the agent’s learning rate *α*), a number of “failures” can be seen.

**Figure 3—figure supplement 2.**
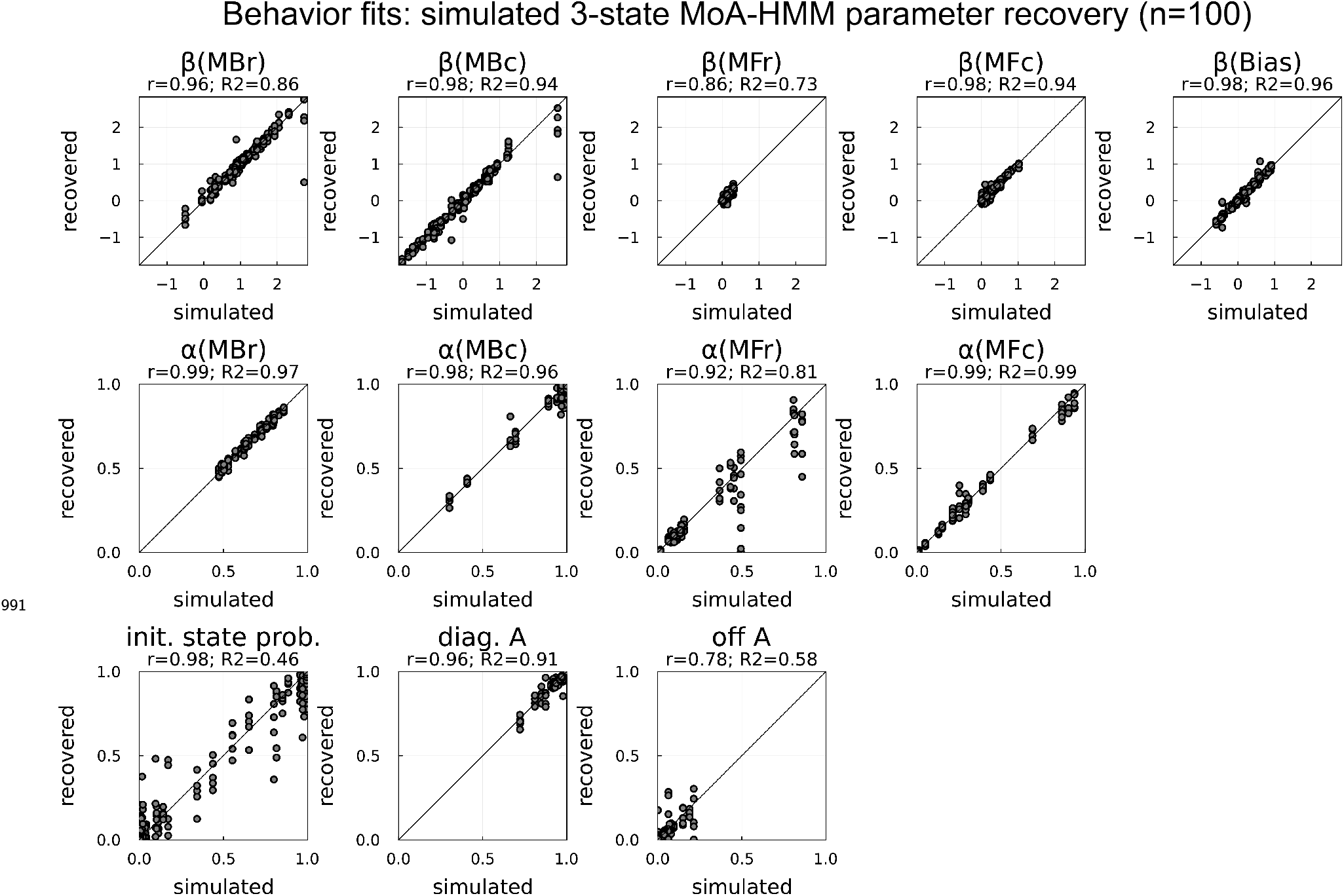
3-state MoA-HMM parameter recovery from data simulated from each rats’ behavioral model fit. Each rat’s model (n=20) was used to generate 5 independent data sets, where each data set contained the same number of trials and sessions as the corresponding rat’s behavioral data set used to fit the behavioral model, giving a total of 100 simulations. “r” is the Pearson’s correlation coefficient between the simulated and recovered parameters, and “R2” is the coefficient of determination, *R*^2^, calculating how well the simulated parameters predict the recovered parameters. Parameters inferred from real data are more reliably recovered.

**Figure 4—figure supplement 1.**
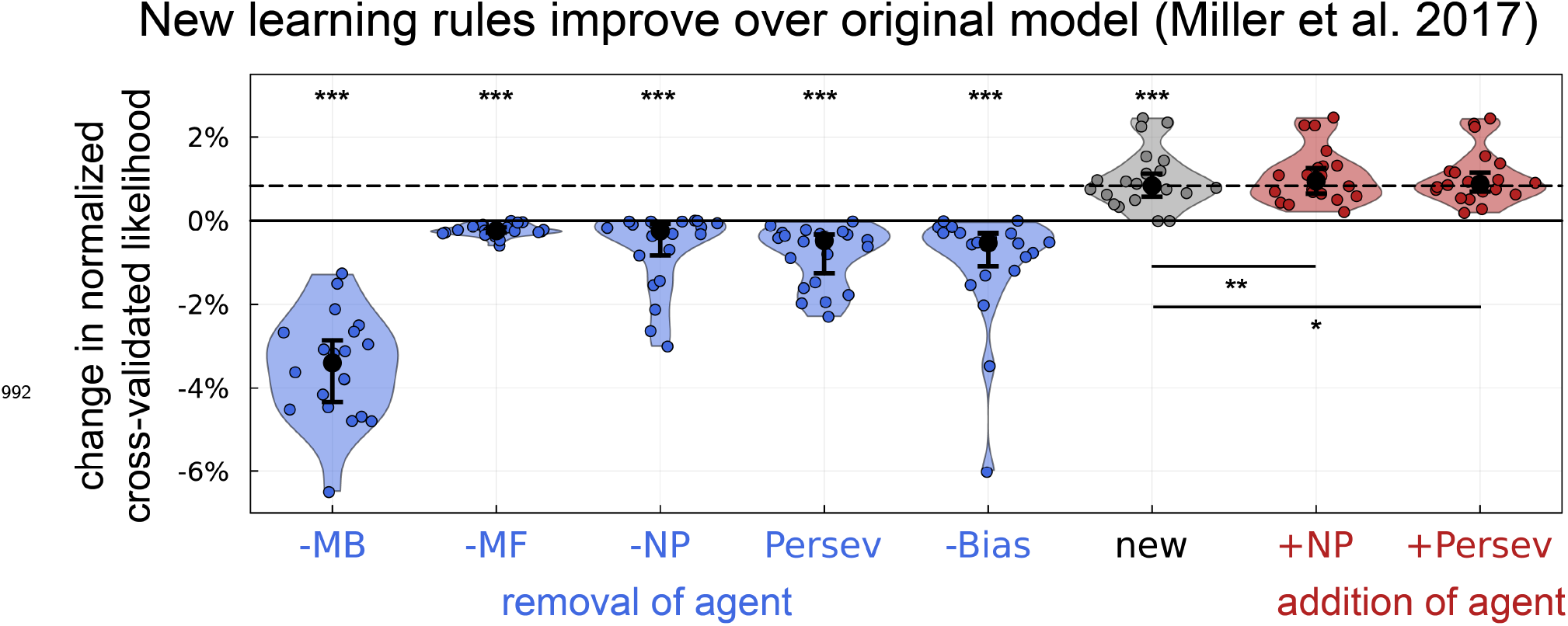
New RL learning rules significantly improve fit to behavior and capture much of the variance explained by the Novelty Preverence and Perseveration agents of the original model from ***Miller et al. (2017***). *:p<0.02, **:p<0.001, ***:p<1E-4

**Figure 5—figure supplement 1.**
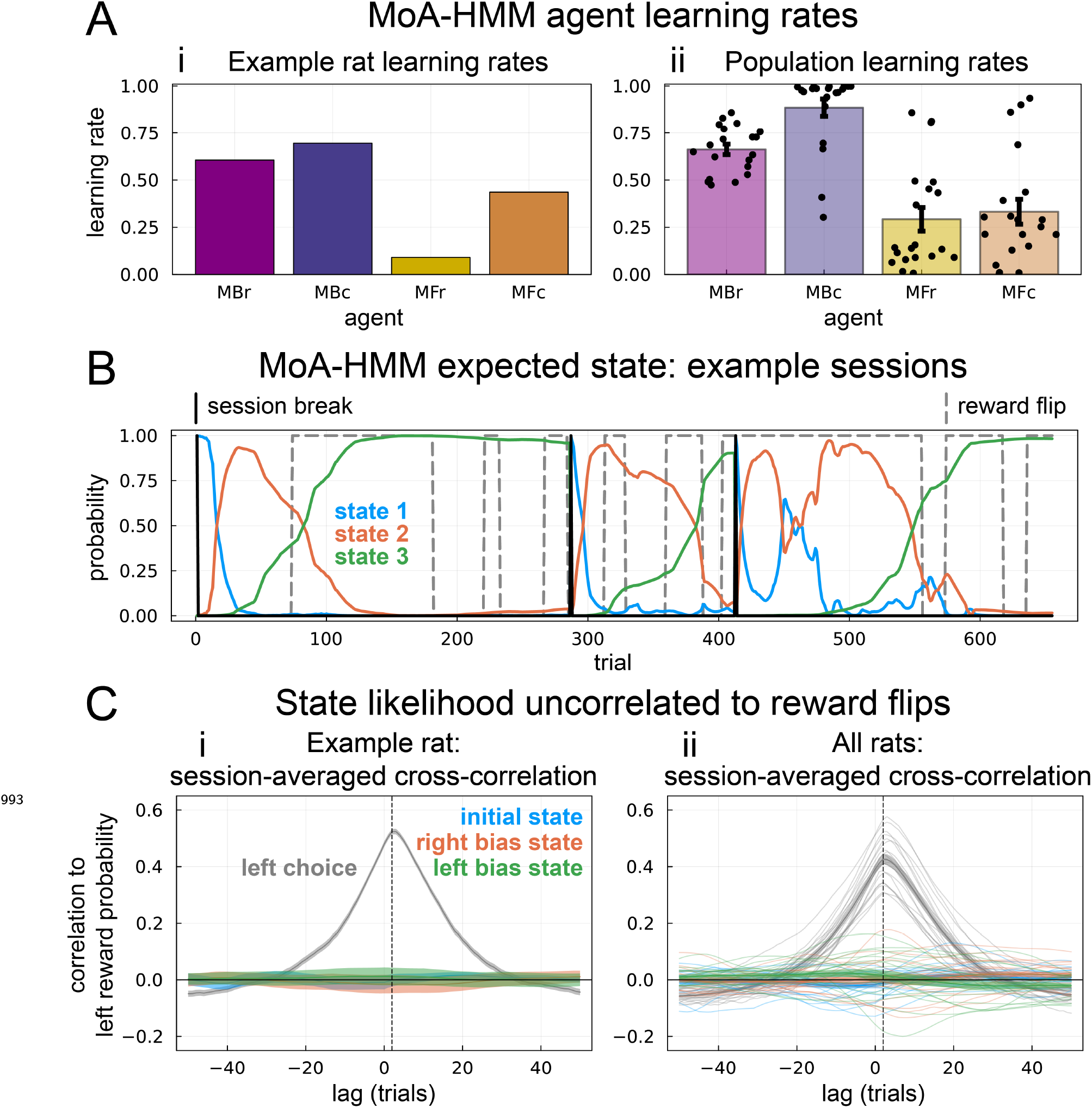
(**A**) Learning rates fit for each agent (**i**) corresponding to the example rat shown in Figure 5A and (**ii**) summarizing each learning rate over the population of rats. Each dot is an individual rat, bars represent the median, and errorbars are bootstrapped 95% confidence intervals around the median. (**B**) Three example sessions showing the inferred state likelihood on each trial from the example rat shown in Figure 5A. (**C**) Cross-correlation between left choices and reward probabilities for the common outcome port given that choice (gray). Left choices are highly correlated to left-outcome reward blocks, with the peak correlation at a slight lag (vertical dashed line) indicating the trial at which the rat detects the reward probability flip. To test whether the latent states track reward flips, the cross correlation is also shown between left-outcome reward probability and the likelihood of each state: initial state (blue), the remaining state with a more rightward choice bias (orange), and the remaining state with a more leftward bias (green). These correspond directly to states 1-3 in the example rat (**i**) whose model is shown in Figure 5A. while other rats had states 2 and 3 assigned according to their individual choice biases.

**Figure 5—figure supplement 2.**
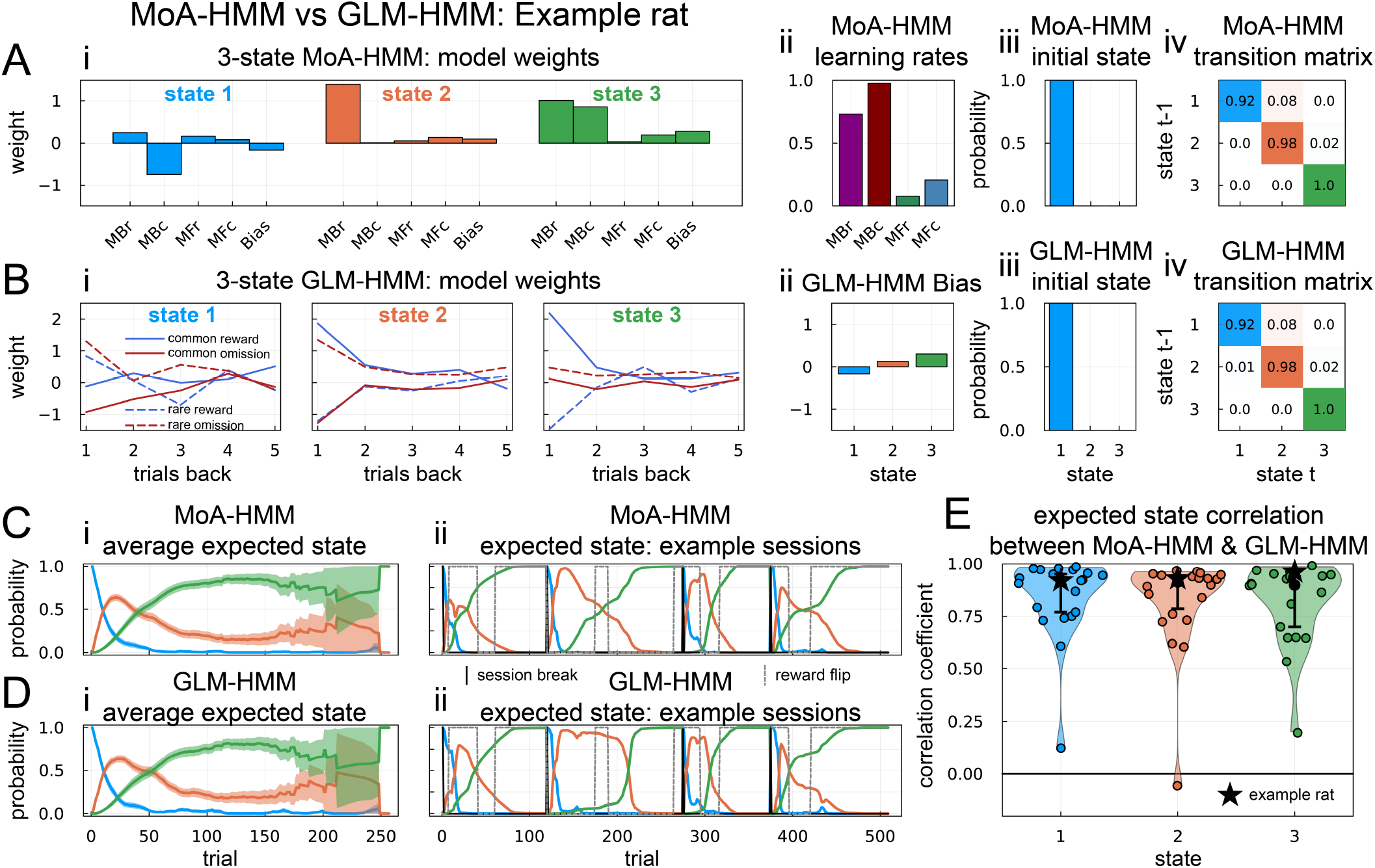
A second 3-state MoA-HMM example and comparison to a GLM-HMM. States identified by a MoA-HMM and GLM-HMM are highly similar. **A** Example 3-state MoA-HMM model parameters with (**i**) agent weights split by state, (**ii**) each agent’s learning rate, (**iii**) the initial state probability, and (**iv**) the state transition matrix. **B** 3-state GLM-HMM fit to the same rat as (A). (**i**) GLM-HMM regression weights for each state. Each state is described by four types of regressors indicating four possible trial types – common-reward, common-omission, rare-reward and rare-omission – and choice direction for up to 5 previous trials, giving 20 parameters per state. Each state additionally had a bias term (**ii**), leading to a total of 63 model weights for a 3-state model. **iii**) GLM-HMM initial state probabilities also identify a prominent initial state. (**iv**) GLM-HMM transition matrix closely matches MoA-HMM. **C** Expected state probabilities for MoA-HMM (**i**) averaged across all sessions and (**ii**) highlighted example sessions. **D** Expected state probabilities for GLM-HMM (**i**) averaged across all sessions and (**ii**) highlighted example sessions. Temporal structure is highly similar to MoA-HMM. **E** Cross-correlation between the expected state probabilities inferred from the MoA-HMM and GLM-HMM (i.e. panels Cii and Dii) across all sessions. Each dot is an individual rat, black circles are medians and error bars are 95% confidence intervals around the median.

**Figure 5—figure supplement 3.**
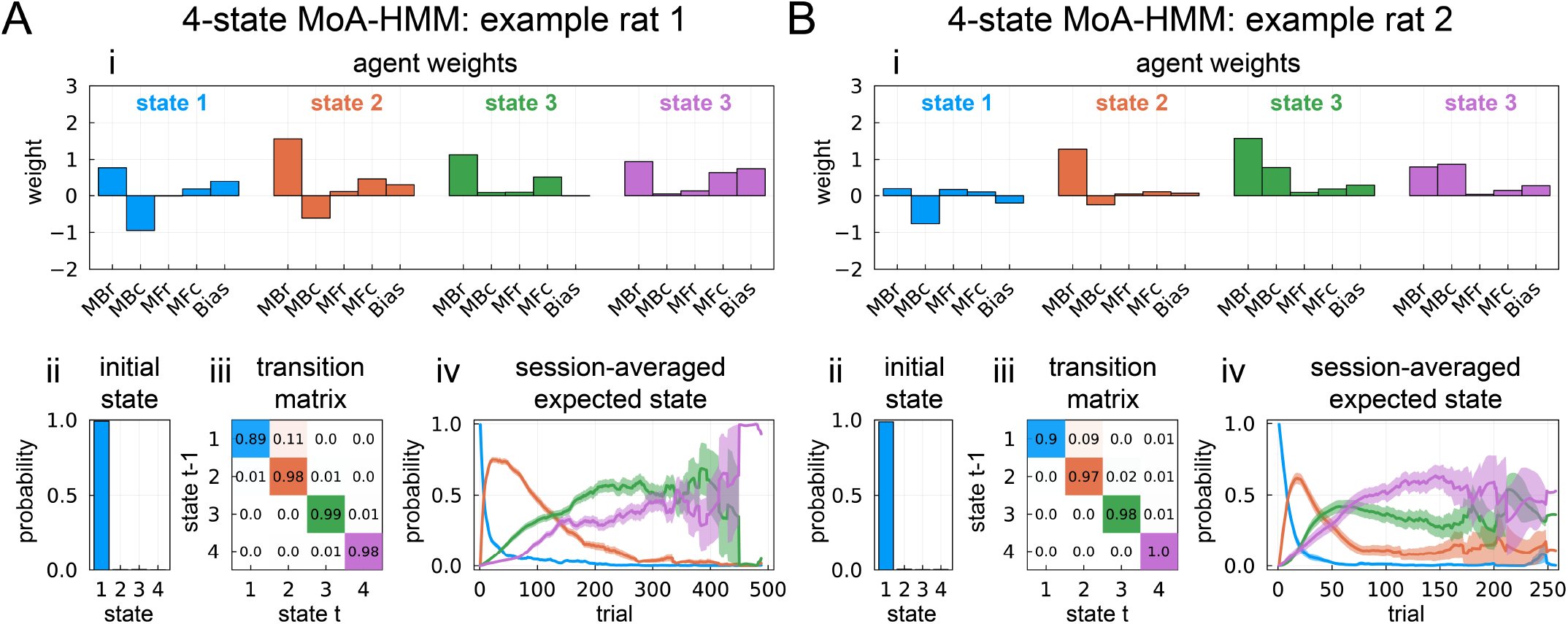
Two example 4-state MoA-HMM fits corresponding to 3 state fits from (**A**) Figure 5A and (**B**) Figure 5-figure supplement 2. States are ordered according to the initial state probability (Aii and Bii) and the transition probabilities to most-likely states that follow (Aiii and Biii). Initial states are generally consistent with the 3-state fits, and the way the remaining two states split into three states is more idiosyncratic. For example, (**A**) suggests state 3 from the smaller model is split into two states (iv) that differ by bias (i), while (**B**) suggest the additional state 4 draws from both the smaller model’s states 2 and 3 (iv), and the state with largest MBr state no longer directly follows the initial state (i).

**Figure 7—figure supplement 1.**
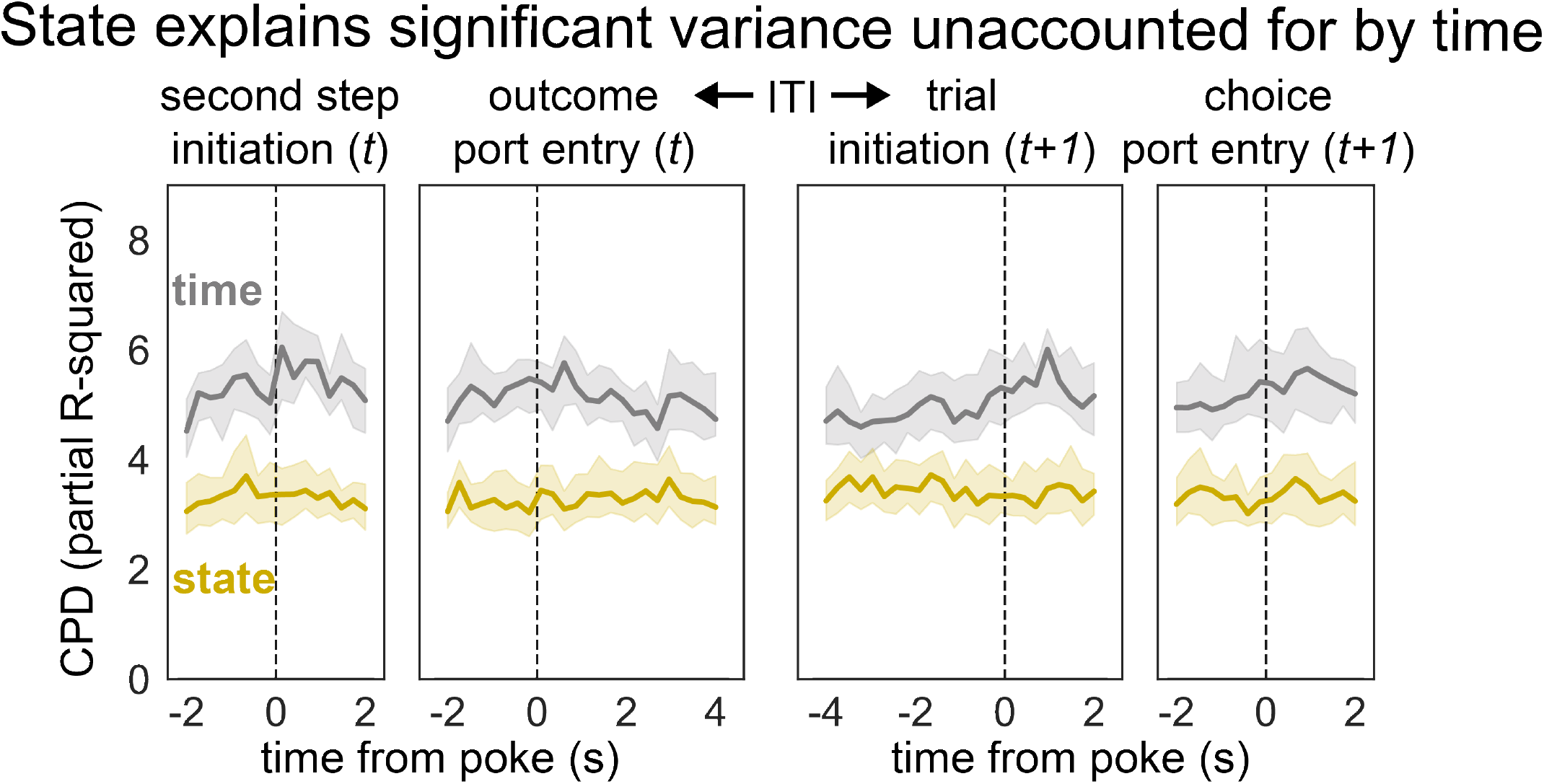
Population CPD computed around the inter-trial interval (ITI) reveals significant encoding of state (gold) even after accounting for time (gray). CPDs were measured as the median CPD across all units, with errorbars corresponding to bootstrapped 95% confidence intervals.

**Figure 7—figure supplement 2.**
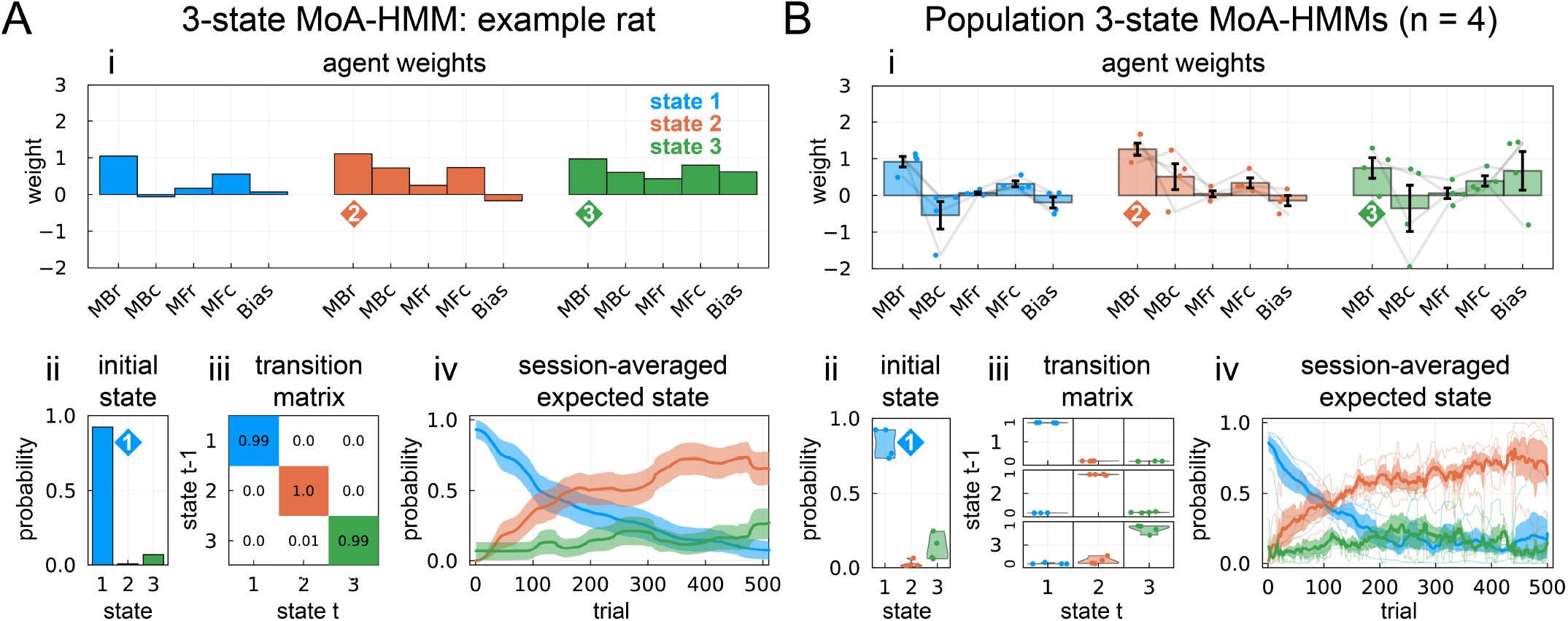
Example and summary 3-state MoA-HMM fits for rats with electrophysiological recordings in OFC.

## References

Akam T, Costa R, Dayan P. Simple Plans or Sophisticated Habits? State, Transition and Learning Interactions in the Two-Step Task. PLOS Computational Biology. 2015 Dec8; 11(12):e1004648. doi: 10.1371/journal.pcbi.1004648.

Akam T, Rodrigues-Vaz I, Marcelo I, Zhang X, Pereira M, Oliveira RF, Dayan P, Costa RM. The Anterior Cingulate Cortex Predicts Future States to Mediate Model-Based Action Selection. Neuron. 2021 Jan; 109(1):149–163.e7. doi: 10.1016/j.neuron.2020.10.013.

Ashwood ZC, Roy NA, Bak JH. Inferring Learning Rules from Animal Decision-Making. NeurIPS. 2020; p. 12.

Ashwood ZC, Roy NA, Stone IR, Urai AE, Churchland AK, Pouget A, Pillow JW. Mice Alternate between Discrete Strategies during Perceptual Decision-Making. Nature Neuroscience. 2022 Feb; 25(2):201–212. doi: 10.1038/s41593-021-01007-z.

Averbeck BB. Theory of Choice in Bandit, Information Sampling and Foraging Tasks. PLOS Computational Biology. 2015 Mar; 11(3):e1004164. doi: 10.1371/journal.pcbi.1004164.

Bishop CM. Pattern Recognition and Machine Learning. Information Science and Statistics, New York: Springer; 2006.

Blanchard TC, Gershman SJ. Pure Correlates of Exploration and Exploitation in the Human Brain. Cognitive, Affective, & Behavioral Neuroscience. 2018 Feb; 18(1):117–126. doi: 10.3758/s13415-017-0556-2.

Bolkan SS, Stone IR, Pinto L, Ashwood ZC, Iravedra Garcia JM, Herman AL, Singh P, Bandi A, Cox J, Zimmerman CA, Cho JR, Engelhard B, Pillow JW, Witten IB. Opponent Control of Behavior by Dorsomedial Striatal Path-ways Depends on Task Demands and Internal State. Nature Neuroscience. 2022 Mar; 25(3):345–357. doi: 10.1038/s41593-022-01021-9.

Calhoun AJ, Pillow JW, Murthy M. Unsupervised Identification of the Internal States That Shape Natural Behavior. Nature Neuroscience. 2019; 22:28.

Camerer C, Ho TH. Experienced-Weighted Attraction Learning in Normal Form Games. Econometrica. 1999; 67(4):827–874.

Cinotti F, Fresno V, Aklil N, Coutureau E, Girard B, Marchand AR, Khamassi M. Dopamine Blockade Impairs the Exploration-Exploitation Trade-off in Rats. Scientific Reports. 2019 May; 9(1):6770. doi: 10.1038/s41598-019-43245-z.

Cockburn J, Man V, Cunningham WA, O’Doherty JP. Novelty and Uncertainty Regulate the Balance between Exploration and Exploitation through Distinct Mechanisms in the Human Brain. Neuron. 2022 Aug; 110(16):2691–2702.e8. doi: 10.1016/j.neuron.2022.05.025.

Correa CG, Griffiths TL, Daw ND, Program-Based Strategy Induction for Reinforcement Learning. arXiv; 2024. doi: 10.48550/arXiv.2402.16668.

Costa VD, Mitz AR, Averbeck BB. Subcortical Substrates of Explore-Exploit Decisions in Primates. Neuron. 2019 Aug; 103(3):533–545.e5. doi: 10.1016/j.neuron.2019.05.017.

Daw ND. Trial-by-Trial Data Analysis Using Computational Models. In: Delgado MR, Phelps EA, Robbins TW, editors. Decision Making, Affect, and Learning: Attention and Performance XXIII Oxford University Press; 2011. p. 0. doi: 10.1093/acprof:oso/9780199600434.003.0001.

Daw ND, Gershman SJ, Seymour B, Dayan P, Dolan RJ. Model-Based Influences on Humans’ Choices and Striatal Prediction Errors. Neuron. 2011 Mar; 69(6):1204–1215. doi: 10.1016/j.neuron.2011.02.027.

Daw ND, Niv Y, Dayan P. Uncertainty-Based Competition between Prefrontal and Dorsolateral Striatal Systems for Behavioral Control. Nature Neuroscience. 2005 Dec; 8(12):1704–1711. doi: 10.1038/nn1560.

Daw ND, O’Doherty JP, Dayan P, Seymour B, Dolan RJ. Cortical Substrates for Exploratory Decisions in Humans. Nature. 2006 Jun; 441(7095):876–879. doi: 10.1038/nature04766.

Dayan P, Daw ND. Decision Theory, Reinforcement Learning, and the Brain. Cognitive, Affective, & Behavioral Neuroscience. 2008 Dec; 8(4):429–453. doi: 10.3758/CABN.8.4.429.

DePasquale B, Brody CD, Pillow JW, Neural Population Dynamics Underlying Evidence Accumulation in Multiple Rat Brain Regions. bioRxiv; 2022. doi: 10.1101/2021.10.28.465122.

Ebitz RB, Albarran E, Moore T. Exploration Disrupts Choice-Predictive Signals and Alters Dynamics in Prefrontal Cortex. Neuron. 2018 Jan; 97(2):450–461.e9. doi: 10.1016/j.neuron.2017.12.007.

Eckstein MK, Collins AG. How the Mind Creates Structure: Hierarchical Learning of Action Sequences. Proceedings of the Annual Meeting of the Cognitive Science Society. 2021; 43(43).

Feher da Silva C, Hare TA. Humans Primarily Use Model-Based Inference in the Two-Stage Task. Nature Human Behaviour. 2020 Oct; 4(10):1053–1066. doi: 10.1038/s41562-020-0905-y.

Gershman SJ. Deconstructing the Human Algorithms for Exploration. Cognition. 2018 Apr; 173:34–42. doi: 10.1016/j.cognition.2017.12.014.

Gillan CM, Kosinski M, Whelan R, Phelps EA, Daw ND. Characterizing a Psychiatric Symptom Dimension Related to Deficits in Goal-Directed Control. eLife. 2016 Mar; 5:e11305. doi: 10.7554/eLife.11305.

Gittins JC. Bandit Processes and Dynamic Allocation Indices. Journal of the Royal Statistical Society Series B (Methodological). 1979; 41(2):148–177.

Gremel CM, Costa RM. Orbitofrontal and Striatal Circuits Dynamically Encode the Shift between Goal-Directed and Habitual Actions. Nature Communications. 2013 Aug; 4(1):2264. doi: 10.1038/ncomms3264.

Groman SM, Massi B, Mathias SR, Curry DW, Lee D, Taylor JR. Neurochemical and Behavioral Dissections of Decision-Making in a Rodent Multistage Task. Journal of Neuroscience. 2019 Jan; 39(2):295–306. doi: 10.1523/JNEUROSCI.2219-18.2018.

Groman SM, Massi B, Mathias SR, Lee D, Taylor JR. Model-Free and Model-Based Influences in Addiction-Related Behaviors. Biological Psychiatry. 2019 Jun; 85(11):936–945. doi: 10.1016/j.biopsych.2018.12.017.

Hasz BM, Redish AD. Deliberation and Procedural Automation on a Two-Step Task for Rats. Frontiers in Integrative Neuroscience. 2018 Aug; 12:30. doi: 10.3389/fnint.2018.00030.

Hogeveen J, Mullins TS, Romero JD, Eversole E, Rogge-Obando K, Mayer AR, Costa VD. The Neurocom-putational Bases of Explore-Exploit Decision-Making. Neuron. 2022 Jun; 110(11):1869–1879.e5. doi: 10.1016/j.neuron.2022.03.014.

Huys QJM, Eshel N, O’Nions E, Sheridan L, Dayan P, Roiser JP. Bonsai Trees in Your Head: How the Pavlovian System Sculpts Goal-Directed Choices by Pruning Decision Trees. PLOS Computational Biology. 2012 Mar; 8(3):e1002410. doi: 10.1371/journal.pcbi.1002410.

Ito M, Doya K. Validation of Decision-Making Models and Analysis of Decision Variables in the Rat Basal Ganglia. Journal of Neuroscience. 2009 Aug; 29(31):9861–9874. doi: 10.1523/JNEUROSCI.6157-08.2009.

Ji-An L, Benna MK, Mattar MG, Automatic Discovery of Cognitive Strategies with Tiny Recurrent Neural Networks. bioRxiv; 2023. doi: 10.1101/2023.04.12.536629.

Killcross S, Coutureau E. Coordination of Actions and Habits in the Medial Prefrontal Cortex of Rats. Cerebral Cortex. 2003 Apr; 13(4):400–408. doi: 10.1093/cercor/13.4.400.

Kool W, Cushman FA, Gershman SJ. When Does Model-Based Control Pay Off? PLOS Computational Biology. 2016 Aug; 12(8):e1005090. doi: 10.1371/journal.pcbi.1005090.

Krueger PM, Wilson RC, Cohen JD. Strategies for Exploration in the Domain of Losses. Judgment and Decision Making. 2017 Mar; 12(2):104–117. doi: 10.1017/S1930297500005659.

Le NM, Yildirim M, Wang Y, Sugihara H, Jazayeri M, Sur M. Mixtures of Strategies Underlie Rodent Behavior during Reversal Learning. PLOS Computational Biology. 2023 Sep; 19(9):e1011430. doi: 10.1371/journal.pcbi.1011430.

Lee SW, Shimojo S, O’Doherty JP. Neural Computations Underlying Arbitration between Model-Based and Model-free Learning. Neuron. 2014 Feb; 81(3):687–699. doi: 10.1016/j.neuron.2013.11.028.

Li JJ, Shi C, Li L, Collins AGE. Dynamic Noise Estimation: A Generalized Method for Modeling Noise Fluctuations in Decision-Making. Journal of Mathematical Psychology. 2024 Apr; 119:102842. doi: 10.1016/j.jmp.2024.102842.

Luo TZ, Kim TD, Gupta D, Bondy AG, Kopec CD, Elliot VA, DePasquale B, Brody CD, Transitions in Dynamical Regime and Neural Mode Underlie Perceptual Decision-Making. bioRxiv; 2023. doi: 10.1101/2023.10.15.562427.

Miller KJ, Botvinick MM, Brody CD. Dorsal Hippocampus Contributes to Model-Based Planning. Nature Neuro-science. 2017 Sep; 20(9):1269–1276. doi: 10.1038/nn.4613.

Miller KJ, Botvinick MM, Brody CD. Value Representations in the Rodent Orbitofrontal Cortex Drive Learning, Not Choice. eLife. 2022 Aug; 11:e64575. doi: 10.7554/eLife.64575.

Miller KJ, Brody CD, Botvinick MM. Identifying Model-Based and Model-Free Patterns in Behavior on Multi-Step Tasks. Neuroscience; 2016.

Miller KJ, Eckstein M, Botvinick MM, Kurth-Nelson Z, Cognitive Model Discovery via Disentangled RNNs. bioRxiv; 2023. doi: 10.1101/2023.06.23.546250.

Miller KJ, Shenhav A, Ludvig EA. Habits without Values. Psychological Review. 2019 Mar; 126(2):292–311. doi: 10.1037/rev0000120.

O’Doherty JP, Lee SW, Tadayonnejad R, Cockburn J, Iigaya K, Charpentier CJ. Why and How the Brain Weights Contributions from a Mixture of Experts. Neuroscience & Biobehavioral Reviews. 2021 Apr; 123:14–23. doi: 10.1016/j.neubiorev.2020.10.022.

Oostland M, Kislin M, Chen Y, Chen T, Venditto SJ, Deverett B, Wang SSH, Cerebellar Acceleration of Learning in an Evidence-Accumulation Task. bioRxiv; 2022. doi: 10.1101/2021.12.23.474034.

Park H, Lee D, Chey J. Stress Enhances Model-Free Reinforcement Learning Only after Negative Outcome. PLOS ONE. 2017 Jul; 12(7):e0180588. doi: 10.1371/journal.pone.0180588.

Roy NA, Bak JH, Akrami A, Brody CD, Pillow JW. Extracting the Dynamics of Behavior in Sensory Decision-Making Experiments. Neuron. 2021 Feb; 109(4):597–610.e6. doi: 10.1016/j.neuron.2020.12.004.

Russek EM, Momennejad I, Botvinick MM, Gershman SJ, Daw ND. Predictive Representations Can Link Model-Based Reinforcement Learning to Model-Free Mechanisms. PLOS Computational Biology. 2017 Sep; 13(9):e1005768. doi: 10.1371/journal.pcbi.1005768.

Schultz W, Dayan P, Montague PR. A Neural Substrate of Prediction and Reward. Science. 1997 Mar; 275(5306):1593–1599. doi: 10.1126/science.275.5306.1593.

Shahar N, Hauser TU, Moutoussis M, Moran R, Keramati M, Consortium N, Dolan RJ. Improving the Reliability of Model-Based Decision-Making Estimates in the Two-Stage Decision Task with Reaction-Times and Drift-Diffusion Modeling. PLOS Computational Biology. 2019 Feb; 15(2):e1006803. doi: 10.1371/journal.pcbi.1006803.

Sutton RS, Barto A. Reinforcement Learning: An Introduction. Second edition ed. Adaptive Computation and Machine Learning, Cambridge, Massachusetts London, England: The MIT Press; 2020.

Wilson R, Wang S, Sadeghiyeh H, Cohen JD, Deep Exploration as a Unifying Account of Explore-Exploit Behavior. OSF; 2020. doi: 10.31234/osf.io/uj85c.

Wilson RC, Bonawitz E, Costa VD, Ebitz RB. Balancing Exploration and Exploitation with Information and Randomization. Current Opinion in Behavioral Sciences. 2021 Apr; 38:49–56. doi: 10.1016/j.cobeha.2020.10.001.

Wilson RC, Geana A, White JM, Ludvig EA, Cohen JD. Humans Use Directed and Random Exploration to Solve the Explore–Exploit Dilemma. Journal of experimental psychology General. 2014 Dec; 143(6):2074–2081. doi: 10.1037/a0038199.

